# Intraspecific variation in functional strategies underlies susceptibility to a vascular disease in grapevine

**DOI:** 10.64898/2025.12.15.694483

**Authors:** Pierre Gastou, Audrey Morin, Nathalie Ferrer, Lauriane Alazet, Régis Burlett, Sylvain Delzon, Frederic Lens, Samuele Moretti, Claudia Rouveyrol, Pierre Pétriacq, Isabelle Svahn, Chloé E. L. Delmas

## Abstract

Woody plant dieback involves complex interactions between pathogens, plant functional traits and environmental conditions. The role of intraspecific variation in functional strategies in shaping susceptibility to dieback remains unclear, despite its importance for understanding plant adaptation to changing environments.

We used a common garden containing 46 *Vitis vinifera* (grapevine) cultivars to test the hypothesis that differences in grapevine susceptibility to esca — a complex vascular disease leading to dieback — result from intraspecific syndromes of leaf gas exchange, wood anatomical and metabolic traits, and microbiome.

Cultivars with conservative water use strategies tended to be less susceptible to esca whereas xylem anatomy did not affect esca susceptibility. Across cultivars, symptomatic plants displayed decreases in leaf gas exchange, stem starch storage and theoretical hydraulic conductivity in response to esca. Symptomatic stems accumulated more secondary metabolites (mostly glycosylated flavonoids and terpenes) in highly susceptible than in weakly susceptible genotypes, whereas stem microbial communities were unaffected. This suggests that disease expression is linked to cultivar-specific metabolic responses.

Intraspecific variation in physiological strategies contributes to differences in susceptibility to a complex vascular disease. Integrative studies unravelling plant-microorganism- environment interactions are crucial to improve our understanding of complex plant diseases.

## Introduction

Wood performs essential functions, such as mechanical support, the transport of water and nutrients, the storage of biochemical compounds and responses to abiotic and biotic stresses (Chave *et al*., 2009; Torres-Ruiz *et al*., 2024). Xylem physiology therefore makes a major contribution to perennial plant functioning and susceptibility to dieback events. The abiotic factors underlying vascular dysfunction are well documented, but increasing attention is now being paid to woody tissues as critical interfaces between the plant and its biotic environment (Torres-Ruiz *et al*., 2024).

Woody tissues host a complex and specific microbiome (Bettenfeld *et al*., 2020; Hamaoka *et al*., 2022), including vascular pathogens that can colonise xylem tissues and induce hydraulic disorders, with potentially detrimental consequences for plant health and survival (Yadeta & Thomma 2013). The physiological consequences of vascular diseases include decreases in carbon assimilation and hydraulic conductivity, starch depletion, and axial and radial growth (Trapero *et al*., 2018; Fanton *et al*., 2022; Dell’Acqua *et al*., 2024). Vascular diseases can also affect endophytic microbial communities in woody tissues depending on the pathosystem and environment (Raghavendra *et al*., 2017; Steinrucken *et al*., 2016; Del Frari *et al*., 2019; Gastou *et al*., 2026).

Xylem functioning is directly involved in physical and metabolic defences against pests and diseases (Shigo, 1984; Pearce, 1996). Infected plants produce vascular occlusions (i.e. tyloses, gums and gels) that limit the axial movements of pathogens and phytotoxins (Kashyap *et al*., 2020). This mechanism slows infection but also impairs water conduction by the xylem sometimes leading to severe hydraulic and photosynthetic dysfunctions, as demonstrated for esca and Pierce’s disease in grapevine (Bortolami *et al*., 2019; Fanton *et al*., 2022), and *Phytophthora ramorum* infection in tanoak (Collins *et al*., 2009). Woody tissues expressing disease symptoms have been shown to display biochemical adjustments, including: (i) impaired primary metabolism, carbohydrate production and storage (Spagnolo *et al*., 2014; Bortolami *et al*., 2021b), (ii) lignin deposition (Sabella *et al*., 2018; Silva *et al*., 2020), and (iii) enhanced production of secondary metabolites and lipid compounds involved in plant stress responses (Lemaitre-Guillier *et al*., 2020; Khattab *et al*., 2021; Chambard *et al*., 2026).

Finally, wood hydraulic properties can regulate plant-pathogen relationships. Few studies have assessed the impact of hydraulic functional traits on susceptibility to vascular disease, despite suggestions of a key role of transpiration and water use efficiency (Bortolami *et al*., 2021b; Gastou *et al*., 2024). Wood anatomy can also interfere with plant-vascular disease interactions. For instance, traits affecting hydraulic conductivity (e.g. vessel diameter) have been shown to be associated with susceptibility to *Xylella fastidiosa* in olive tree and grapevine (Chatelet *et al*., 2011; Petit *et al*., 2021), *Ophiostoma novo-ulmi* susceptibility in elm (Venturas *et al*., 2013) and *Raffaelea lauricola* susceptibility in avocado (Beier *et al*., 2021). The fine structure of intervessel bordered pits strongly influences elm resistance to *O. novo-ulmi* (Martín *et al*., 2009) but not grapevine susceptibility to Pierce’s disease (Fanton & Brodersen, 2021; Fanton *et al*., 2024). Wood functional traits influence genotypic susceptibility to plant disease either directly, through effects on the spread of pathogens and toxic metabolites, or indirectly, by modulating physiological processes that affect disease susceptibility. Hydraulic properties (i.e. ecophysiological and anatomical traits), in particular, are under moderate-to-strong genetic control and contribute to between and within-species functional adaptation (Lauri *et al*., 2011; Venturas *et al*., 2013; Chhetri *et al*., 2020). Plant responses to vascular disease also vary within species. Tolerant genotypes often display earlier and stronger defence responses, potentially limiting pathogen development and spread (Gomez-Gallego *et al*., 2019; Silva *et al*., 2020). By contrast, highly susceptible genotypes typically suffer more severe physiological disruption, including a loss of hydraulic conductivity due to occlusions, leading to lower levels of carbon assimilation and starch reserves in plant organs (Fanton *et al*., 2022). Despite this observed intraspecific variability, the functional determinants of disease susceptibility across genetically diverse perennial species remain poorly understood. Integrative approaches spanning multiple dimensions of plant functions are therefore needed to understand how variation in plant functional traits and microbial diversity shapes plant responses to chronic biotic stress and, ultimately, resilience under interacting environmental constraints.

Here, we investigated whether intraspecific variation in functional traits explains cultivar differences in susceptibility to esca. Esca is a complex vascular disease that poses a serious threat to cultivated grapevine (*Vitis vinifera* L.), a perennial crop exhibiting considerable genetic and phenotypic diversity (Keller, 2020). Esca is characterised by erratic foliar symptoms developing over the summer (Lecomte *et al*., 2024) depending on cultivar, plant age, and climatic conditions (Etienne *et al*., 2024, 2025), as well as internal wood decay caused by fungal pathogens in the trunk (Bruez *et al*., 2020; Gastou *et al.,* 2026). It induces clear physiological modifications and hydraulic impairment in both woody and vegetative tissues (Bortolami *et al*., 2019; 2021a; Dell’Acqua *et al*., 2024, Chambard *et al*., 2026). Its impact on vine health, yield and fruit quality is jeopardising the long-term sustainability of vineyards worldwide (Dewasme *et al*., 2022). Few cost-effective control methods are currently available, but genetic leverage (i.e. cultivar choice and varietal selection) is a promising tool for the sustainable management of esca as there is a high intraspecific diversity of esca susceptibility among *Vitis vinifera* cultivars (Etienne *et al*., 2024; Gastou *et al*., 2024). Esca is, therefore, a relevant case study for explorations of the modulation of perennial plant susceptibility to vascular diseases at genotype level. The physiological response to esca also varies within small ranges of genotypes, including differences in defence responses (e.g. the production of vascular occlusions and secondary metabolites; Lemaitre-Guillier *et al*., 2020; Bortolami *et al*., 2023) or stem physiology (e.g. changes in the timing and sequence of stem radial growth; Dell’Acqua *et al*., 2024). Nevertheless, we still know little about the functional traits underlying grapevine adaptation to esca under natural infection conditions.

We used a common garden planted with up to 46 mature grapevine genotypes (*V. vinifera* L.) with very different levels of susceptibility to esca (Gastou *et al*., 2024). We performed an integrative study to characterise plant hydraulic and wood anatomical traits, carbon assimilation, starch storage, and the metabolic and microbial composition of the wood, in both asymptomatic and esca-symptomatic plants (Fig. 1). We addressed the following questions: Are differences in disease susceptibility associated with coordinated variation in water-use, carbon-use and defence-related traits among cultivars? Does cultivar susceptibility modulate the nature and intensity of plant physiological, microbial metabolic responses to esca?

**Figure 1.**
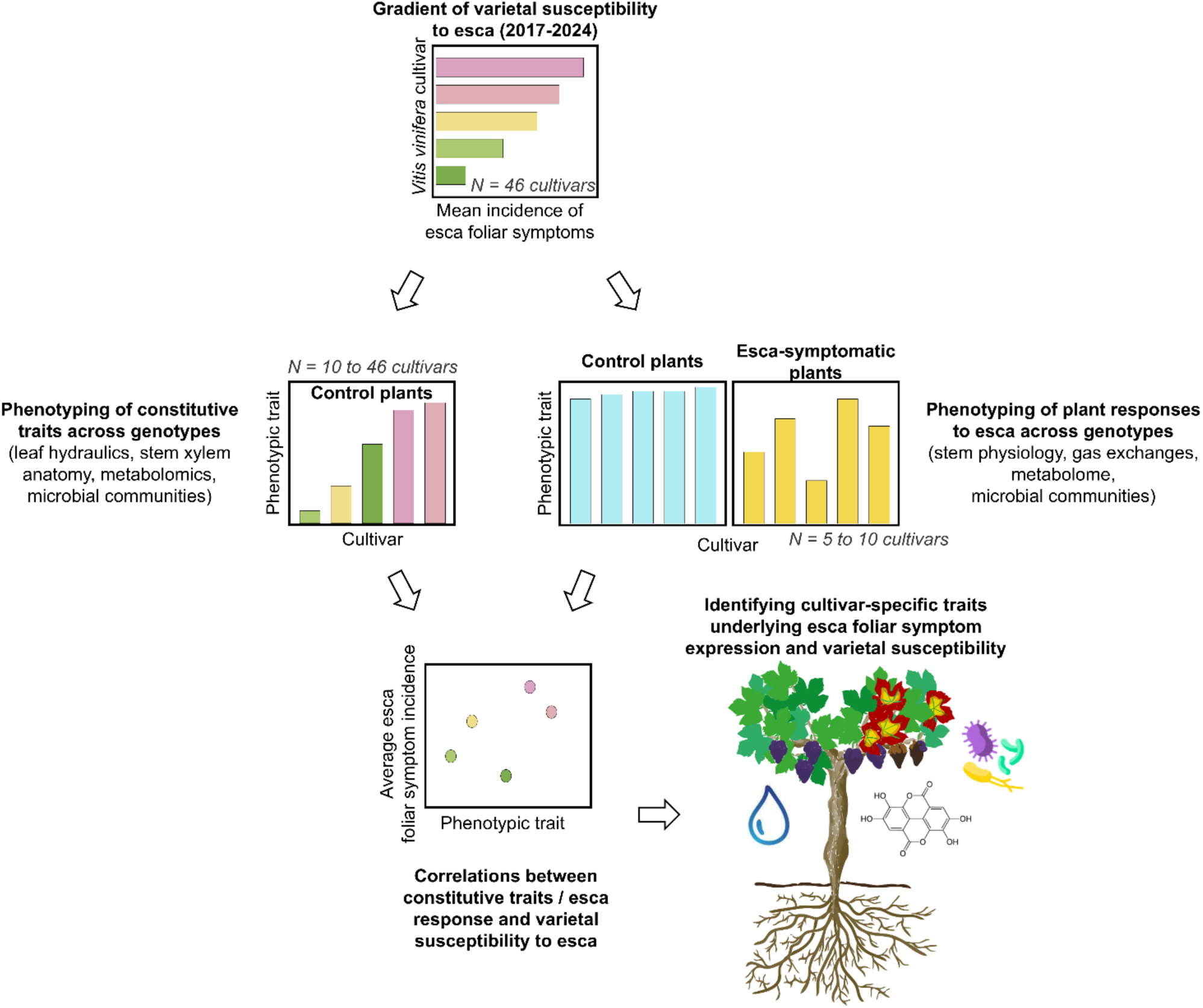
Conceptual framework used to identify functional strategies and microbial drivers of intraspecific differences in esca susceptibility across 46 *Vitis vinifera* cultivars. Esca susceptibility was assessed on the basis of the mean incidence of foliar symptoms from 2017 to 2024. Constitutive traits for each genotype were phenotyped in control plants only. Plant responses to esca were investigated by comparing control and esca-symptomatic plants. We assessed the correlation between each constitutive/response-related trait and esca varietal susceptibility, discussing and ranking these correlations, leading to the development of a preliminary conceptual framework for esca susceptibility. The colours of the bars and dots are coloured indicate varietal susceptibility to esca, from green (weakly susceptible) to red (highly susceptible), or plant responses, with bars are coloured light blue for control plants and yellow for esca-symptomatic plants. These colours are used consistently throughout the article. Mean esca symptom severity in diseased plants was similar across genotypes.

We hypothesised that (i) cultivars expressing acquisitive resource-use strategies (high leaf gas exchange rates and vigorous growth) would be more susceptible to esca than cultivars displaying more conservative strategies; (ii) perturbations in stem primary microbial and metabolism communities would be greater in susceptible cultivars, reflecting lower tolerance to vascular dysfunction; and (iii) plant defence responses to esca involve similar mechanisms in all genotypes but are weaker in susceptible cultivars.

## Materials and methods

### Common garden experimental vineyard and esca foliar symptom monitoring

Plants were grown and sampled in the VitAdapt common garden experimental vineyard at the Institut National de Recherche pour l’Agriculture, l’Alimentation et l’Environnement (INRAE) research station (Villenave d’Ornon, Nouvelle-Aquitaine, France), at 44°47’23.83 N’’, 0°34’39.3’ W’. We focused on *V. vinifera* L. cultivars (Supplementary Table S1) planted from 2009 to 2010 in a randomized block design, grafted onto Selection Oppenheim 4 (SO4) clone 761, as described in detail in Gastou *et al*. (2024). We monitored these plants (*n* = 1,840, 40 plants per cultivar, 46 cultivars) for esca foliar symptoms and dieback each summer from 2017 to 2023 (Gastou *et al*., 2024), and again in 2024. The incidence and severity of esca foliar symptoms (classified as light *vs.* intermediate *vs.* severe at the canopy level as described by Gastou *et al*., 2024) were recorded at individual plant level. We calculated varietal esca foliar symptom incidence — i.e. mean incidence of foliar symptoms by cultivar between 2017 and 2024, hereafter referred to as varietal mean esca susceptibility — and analysed its gradient. K-means clustering was also performed on the varietal incidence of esca foliar symptoms, with three clusters and 10 random initialisations used to create three esca varietal susceptibility classes in some analyses: low (25 cultivars), moderate (12 cultivars) and high (9 cultivars) (Supplementary Table S1). For subsequent experiments, cultivars were distributed along this gradient and across these three clusters.

The severity of esca foliar symptoms was not correlated with incidence at cultivar level (Gastou *et al*., 2024). Diseased plants of cultivars for which the incidence of symptomatic plants was low could, therefore, nevertheless display severe symptoms, and the converse was also true. None of the plants in this vineyard was artificially inoculated with wood pathogens and dead vines were regularly removed to prevent local hotspots of spore production. We can therefore assume that all plants were exposed to similar risks of natural pathogen infection. All the stem samples studied were collected from three of the four experimental blocks of the common garden vineyard with similar incidences of esca foliar symptoms (blocks 1, 3 and 5; as presented by Gastou *et al*., 2024).

### Gas exchange measurements

On 12^th^ and 13^th^ June 2024, we measured stomatal conductance under natural conditions on all plants of the VitAdapt vineyard (*n* = 1,840, 40 plants per cultivar, 46 cultivars). Stomatal conductance g_s_ (mmol m^−2^s^−1^) was measured between 9 and 11 am, with a porometer (LI- 600, LI-COR, Lincoln, USA; Table 1). During the performance of these measurements, photosynthetic radiation ranged from 1600 to 1890 µmol m^-2^ s^-1^, with a temperature of about 21°C and a relative humidity of about 40%. The mean vapour pressure deficit (VPD) was 1.6 ± 0.2 kPa.

**Table 1.** Overall statistics for hydraulic, anatomical and physiological traits measured in the VitAdapt common garden vineyard (Villenave d’Ornon, France) . Information includes the measurements and calculation formulae, number of observations, mean values, standard deviations, relative standard deviations, and varietal minimum - maximum values. A zone corresponds to three vascular bundles in either the dorsoventral or lateral region of the stem

| Trait | Abbreviation | Measurement - Formula | Cultivars (replicates) | mean ± SD | RSD | min | max |
| --- | --- | --- | --- | --- | --- | --- | --- |
| Stomatal conductance<br>(mmol m<sup>-2</sup> s<sup>-1</sup>) | g<sub>s</sub> | 1 green leaf per plant | 46 cultivars<br>(n = 730 leaves) | 167.2 ± 88.6 | 53.0% | 97.6<br>(Cot) | 249.4<br>(Arinarnoa) |
| Minimum leaf conductance 6h-12h<br>(mmol m<sup>-2</sup> s<sup>-1</sup>) | g<sub>min</sub> | 1 green leaf per plant | 19 cultivars<br>(n = 113 leaves) | 7.63 ± 1.07 | 14.0% | 5.94<br>(Cabernet Franc) | 9.66<br>(Tempranillo) |
| Specific theoretical hydraulic conductivity<br>(kg s<sup>-1</sup> m<sup>-1</sup> MPa<sup>-1</sup>)<br>[2 zones per stem, D<sub>vessels</sub> > 25µm] | Specific K<sub>th</sub> | $K_{th} = \frac{\sum(\frac{\pi * \Phi^4 * \rho}{128 * \eta})}{A}$ | 10 cultivars<br>(n = 120 zones) | 70.3 ± 59.4 | 84.5% | 41.7<br>(Gamay) | 120.5<br>(Sauvignon Blanc) |
| Normalised percent of xylem ray parenchyma<br>[2 zones per stem] | F<sub>RP</sub> | $\frac{ray\ parenchyma\ surface\ area}{total\ xylem\ surface\ area}$ | 10 cultivars<br>(n = 120 zones) | 13.4 ± 3.5 | 26.0% | 9.9<br>(Gamay) | 20.4<br>(Chasselas) |
| Aperture fraction of intervessels pits (%)<br>[several zones per stem] | F<sub>AP</sub> | $\frac{pit\ aperture\ surface\ area}{total\ pit\ surface\ area}$ | 13 cultivars<br>(n = 3076 pits) | 12.9 ± 3.8 | 29.4% | 10.3<br>(Merlot) | 15.9<br>(Chardonnay) |
| Xylem ray parenchyma layers | | 2 zones per stem | 10 cultivars<br>(n = 120 zones) | 5.1 ± 1.1 | 20.1% | 4.3<br>(Gamay) | 7.2<br>(Chasselas) |
| Xylem fibre lumen area<br>(µm<sup>2</sup>) | A<sub>flumen</sub> | 4 zones per stem, 25 fibres / zone | 10 cultivars<br>(n = 240 zones) | 170.7 ± 27.4 | 16.0% | 137.5<br>(Chasselas) | 195.9<br>(Sauvignon Blanc) |
| Intervessel pit density<br>(µm<sup>-1</sup>)<br>[several zones per stem] | N<sub>p</sub> | $\frac{pit\ number}{vessel\ element\ length}$ | 13 cultivars<br>(n = 236 zones) | 0.18 ± 0.03 | 14.3% | 0.16<br>(Cabernet-Sauvignon) | 0.19<br>(Chasselas) |
| Percent loss of theoretical hydraulic conductivity driven by vascular occlusion<br>[2 zones per esca-symptomatic stem, D<sub>vessels</sub> > 25µm] | Occlusion<br>PLC<sub>th</sub> | $\frac{K_{th}\ of\ occluded\ vessels}{Total\ K_{th}}$ | 9 cultivars<br>(n = 100 zones) | 22.1 ± 29.3 | 132.7% | 11.7<br>(Chenin) | 38.8<br>(Ugni Blanc) |
| Percentage of ray parenchyma area storing starch [control stems, 20 rays per stem] | P<sub>starch</sub> | $\frac{\text{black} - \text{colored ray parenchyma area}}{\text{total ray parenchyma surface area}}$ | 10 cultivars<br>(n = 720 rays) | 83.7 ± 13.5 | 16.1% | 66.7<br>(Chenin) | 95.0<br>(Merlot) |
| Percentage of ray parenchyma area storing starch [esca stems, 20 rays per stem] | P<sub>starch</sub> | $\frac{\text{black} - \text{colored ray parenchyma area}}{\text{total ray parenchyma surface area}}$ | 9 cultivars<br>(n = 500 rays) | 51.3 ± 34.6 | 67.5% | 27.5<br>(Syrah) | 84.1<br>(Chasselas) |
| Percent occluded vessels (%) [esca stems] | | $\frac{\text{Number of occluded vessels}}{\text{Total number of vessels}}$ | 32 cultivars<br>(n = 324 zones) | 21.1 ± 23.5 | 111.7% | 0<br>(Carmenere) | 56.9%<br>(Grenache) |

During the summer of 2024, we performed two campaigns of gas exchange measurements with a portable photosynthesis system (TARGAS-1, PP Systems, Amesbury, USA). The first campaign was performed before esca symptom expression (1^st^ July 2024; 34 plants, 5 cultivars) and the second was performed during the period of symptom expression (2^nd^ August; 37 plants, 5 cultivars). Measurements were taken under clear sky conditions, between 9 am and 11 am (local time), on green leaves from control stems and green leaves from esca/tiger-stripe stems. Maximal CO_2_ assimilation *A*_max_ (µmol m^−2^s^−1^), and maximal stomatal conductance g_s,max_ (mmol m^−2^ s^−1^) were recorded under optimal conditions of photosynthetically active radiation (1500 μmol m^−2^ s^−1^; Table 1). All gas exchange measurements were performed by clamping one healthy, mature, well-exposed leaf per plant.

### Determination of minimum leaf conductance

For each cultivar, water loss from six individual detached leaves was measured with a custom-built system based on the DroughtBox (Billon *et al*., 2020, Burlett *et al*., 2025). The setup, housed in a 1200 L growth chamber (Fitoclima 1200, Aralab, Portugal), used 24 micro-load cells (0–100 g range; 3139_0, Phidgets Inc.) connected via a Wheatstone bridge board (1046_OB, Phidgets Inc.) to record mass loss continuously, at five-minute intervals. Data acquisition and calibration were managed with custom software (Cuticular v1, University of Bordeaux). The chamber was maintained at 25°C and 60% RH (VPD ≈ 1.26 kPa), with top and bottom lighting at about 400 μmol m^−2^ s^−1^. The petiole bases were sealed with paraffin wax to prevent desiccation. Before measurements, leaves were rehydrated in the dark for 8 h and leaf area (A_leaf_) was measured on scanned images (v850 Pro, Epson, Japan) with dedicated software (Winfolia, Regent instrument, Canada). Turgid weight was measured with a four-digit balance (Pioneer, Ohaus, USA). The leaves were dehydrated and oven-dried at 65°C for 72 h to determine dry weight. Minimum conductance was calculated from the rate of water loss (dw/dt) over a six-hour period (from 6 h to 12 h after the start of measurement, i.e. approximately the relative water content in the 50%-70% interval), and normalized by VPD, as shown in Eq 1. This process was automated with custom R scripts (g_Residual, University of Bordeaux).

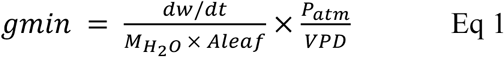

where M_H2O_ is the molecular weight of water (18.01 g mol^-1^), A_leaf_ is leaf area (in m^2^), P_atm_ is the atmospheric pressure in the chamber (c. 101.9 kPa) and VPD is the leaf-to-air vapour pressure deficit (in kPa).

### Anatomical and functional phenotyping of grapevine stems

#### Stem sampling

We studied the impact of cultivar and esca expression on woody stem anatomy and function by sampling a total of 114 current-year stem internodes from individual plants from 33 of the 46 cultivars between 8^th^ and 21^st^ August 2023. In detail, we sampled 35 internodes from control asymptomatic plants (three to six plants per cultivar; 10 cultivars) and 81 internodes from esca/tiger-stripe stems (i.e. esca-symptomatic stems; one to three plants per cultivar; 32 cultivars, including nine from which control stems were also sampled). The seventh internode from the base of a stem of the year concerned was selected from each plant. Samples were stored in 70% ethanol [v/v] for subsequent histological analysis.

#### Histological phenotyping of anatomical and functional xylem traits in stems

For a subsample of 60 internodes (10 cultivars: three weakly susceptible, three moderately susceptible and four highly susceptible; Supplementary Table S1), 15 µm thick cross-sections were cut with a WSL-Lab microtome (modified Reichert-type; WSL Institute, Birmensdorf, Switzerland; Gärtner *et al*., 2015) for the precise visualisation of fibres and parenchyma. Cross-sections were stored on glass slides in a 1:1 water/glycerol mixture before staining. They were processed according to a slightly modified version of the protocol described by Gärtner and Schweingruber (2013): slides were stained for two minutes with a 1:1 mixture of safranin and Astra Blue, rinsed several times, first with water subjected to reverse-osmosis, then with a graded series of ethanol solutions (70%, 96%, 100% [v/v]). The sections were then impregnated with xylene, mounted in Histolaque LMR on a slide and covered with a coverslip. They were then allowed to dry for at least 14 hours. High-resolution images of one section per sample were taken with a NANOZOOMER 2.0HT (Hamamatsu Photonics, Hamamatsu City, Japan; Fig. 2A, B and C).

**Figure 2.**
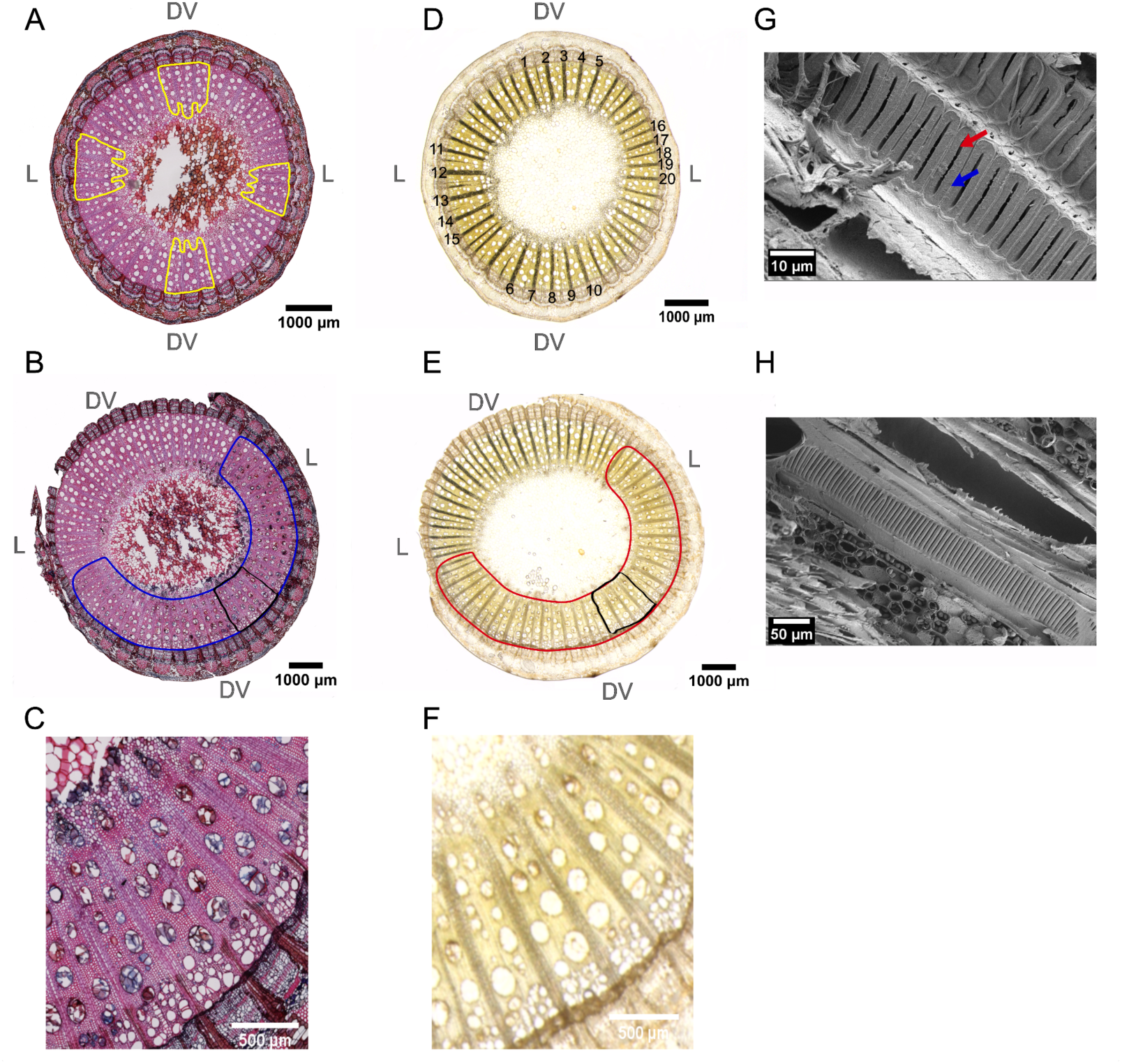
Microscopy images of *Vitis vinifera* stem cross-sections. (A, D) Cross-sections of a control Chenin stem, stained with (A) a mixture of safranin/Astra Blue and (D) Lugol iodine. (B, E) Cross-sections of a Chenin stem displaying severe esca symptoms, stained with (B) a mixture of safranin/Astra Blue, revealing a sector enriched in vascular occlusions (outlined in blue) and (E) Lugol iodine revealing a starch-depleted sector occurring in the same area (outlined in red). (C) Magnified portion of the safranin/Astra Blue-stained cross-section (shown in black on panel B), in which vascular occlusions are clearly visible. (F) Magnified portion of the Lugol iodine-stained cross-section (shown in black on panel E), in which starch- poor parenchyma rays are clearly visible. (G, H) Scanning electron micrographs showing parts of a vessel wall with laterally elongated (scalariform) intervessel pits with visible pit borders (blue arrow) and pit apertures (red arrow), obtained from tangential surfaces of (G) a control Riesling stem and (H) a control Ugni Blanc stem.

We measured various anatomical traits on the images obtained with Image J 1.53t (Wayne Rasband and contributors, National Institutes of Health; https://imagej.net/software/fiji/). In grapevine, the xylem vessels are wider in the dorsoventral part of the stem than in the lateral part of the stem (Brodersen *et al*., 2013; Pouzoulet *et al*. 2020) and they are organised into bundles separated by ray parenchyma (Fig. 2A, B and C). This anatomical diversity of grapevine stems was taken into account by selecting four zones on each cross-section: two zones in the dorsoventral part of the stem and two zones in the lateral part of the stem, each comprising three vascular bundles (Fig. 2A). We filtered xylem vessels on the basis of diameter, retaining only those with a diameter of more than 25 µm, to prevent confusion with tracheids or fibres and for the standardisation of subsequent hydraulic trait calculations. We calculated mean specific theoretical hydraulic conductivity in two zones (specific K_th_), the normalised percentage of xylem ray parenchyma in two zones (i.e. ray parenchyma area over total zone area; F_RP_), the number of xylem ray parenchyma layers measured in the central part of the ray in two zones (number of RP layers), and mean xylem fibre lumen area in four zones (A_flumen_). We also calculated the percentage loss of theoretical hydraulic conductivity due to vascular occlusions as the occluded vessel area over total vessel area in two zones (*n* = 2 zones; occlusion PLC_th_). All measurements and abbreviations are detailed in Table 1.

In another subsample of 54 internodes, 30 µm cross-sections were cut with a GSL-1 microtome (Gärtner *et al*., 2014). These sections were stained and imaged as described above. We determined only the percentage of the total number of vessels that were occluded in four zones (Table 1).

The four study zones for anatomical measurements (each including three vascular bundles) are shown in yellow (panel A). The 20 parenchyma rays considered for starch quantification are numbered in panel D. The dorsoventral part of the stem is indicated as DV and the lateral part as L.

#### Histological quantification of starch in xylem ray parenchyma

For the same subsample of 60 internodes, we visualised starch in the ray parenchyma of xylem cross-sections as described by Fanton *et al*. (2022). Briefly, 30 µm cross-sections were cut with a WSL-Lab microtome, stained by incubation for one minute in a 0.5% iodine tincture obtained by diluting Lugol iodine concentrate (Pro-Lab Diagnostics, Bromborough, United Kingdom). They were then rinsed several times with water treated by reverse-osmosis, mounted on a slide and covered with a coverslip. We immediately photographed one section per sample with a Nikon SMZ1270 camera (Nikon Instruments, Melville, USA) and the NIS-ElementsD software associated with the magnifier (Fig. 2D, E and F).

We used Image J to determine the percentage of the ray parenchyma area storing starch (i.e. iodine-stained amyloplasts). We selected four study zones for each cross-section (Fig. 2D) — two dorsoventral zones and two lateral zones (identified on the basis of vessel size, as described by Pouzoulet *et al*., 2020) — each comprising five parenchyma rays. We then applied a threshold-based segmentation based on the intensity of black pixels. Finally, we quantified the percentage area of each ray in which the intensity of black pixels was above the threshold, as a proxy for the percentage of the ray parenchyma area storing starch (P_starch_; Table 1).

#### Phenotyping of intervessel pits

We assessed the variability in intervessel pit structure between cultivars by sampling the fifth internode from the base of control stems during two summers (three plants per cultivar, 13 cultivars: four weakly susceptible, four moderately susceptible and five highly susceptible; Supplementary Table S1). On 29^th^ August 2023, 18 internodes were sampled from different plants of six cultivars. On 2^nd^ September 2024, 24 internodes were sampled from different plants of eight cultivars (Saperavi was sampled in both these years). Samples were prepared for observation scanning electron microscopy (SEM) according to a protocol adapted from that described by Jansen *et al*. (2008). Tangential planes were split with the WSL-Lab microtome and air-dried at 37°C, without chemical treatment. They were sputter-coated with platinum for 20 s at 15 mA with a Q150T Plus (Quorum Technologies, Lewes, United Kingdom). Observations were performed at an acceleration voltage of 2 kV with a FEG ZEISS Gemini300 high-resolution scanning electron microscope (Carl Zeiss, Oberkochen, Germany), at the Bordeaux Imaging Centre, a member of the national infrastructure France Bio Imaging (Bordeaux, France; ANR-10-INBS-04). On selected high-quality SEM scans (see examples in Fig. 2G and H), for sections of vessels in contact with each other (i.e. with continuous intervessel pitting), we quantified the density of intervessel pits (i.e. number of pits per unit vessel element length; N_P_) and the aperture fraction of the intervessel pits (i.e. ratio of pit aperture area over total pit area; F_AP_; Table 1) with Image J.

### Stem metabolome and microbiome analysis

#### Stem sampling

We investigated the impact of cultivar and esca expression on the stem metabolome and microbial communities by sampling between 8^th^ and 11^th^ August 2023, a total of 88 stem internodes from individual plants from the same 10 cultivars used to study xylem anatomy. The ninth internode from the base of each stem was sampled, collected into a sterile 5 mL microtube and immediately placed in liquid nitrogen. All the equipment was disinfected with 96% ethanol [v/v] between plants. The samples were stored at -80°C for subsequent analyses. We sampled internodes from 29 control stems (three per cultivar, 10 cultivars), 32 esca/asymptomatic stems (i.e. asymptomatic stems from esca-symptomatic plants; three to five per cultivar, eight cultivars) and 27 esca/tiger-stripe stems (two to four per cultivar, nine cultivars; Supplementary Table S1).

#### Metabolite extraction, LC-MS and data processing

Stem samples were ground in liquid nitrogen with a TissueLyser II one-ball mill (Qiagen, Hilden, Germany). Metabolites were extracted from the ground samples, which were freeze-dried with an ALPHA 1-4 LDplus lyophiliser (Martin Christ, Osterode am Harz, Germany). Samples (10 mg) of lyophilised wood tissues were sent to the MetaboHUB-Bordeaux Metabolome facility (https://metabolome.u-bordeaux.fr/; Villenave-d’Ornon, France) for metabolite extraction, LC-MS and preprocessing of the data. Metabolites were extracted automatically with robots as previously described (Dussarrat *et al*., 2022, Luna *et al*., 2020), in a solvent containing ethanol 80% [v/v] and 0.1% formic acid [v/v], with methyl vanillate (250 µg/mL) as an internal standard. Extracts were filtered through MultiScreen GV sterile filtration plates with 0.22 µm pores (Merck, Molsheim, France). Quality controls were prepared by mixing all samples and were included on each plate. Extraction blanks were also included on each plate. Untargeted metabolic profiling by UHPLC-LTQ-Orbitrap mass spectrometry (LCMS) was performed with an Ultimate 3000 ultra-high pressure liquid chromatography (UHPLC) system coupled to an LTQ-Orbitrap Elite mass spectrometer interfaced with an electrospray ionisation source (ESI, Thermo Fisher Scientific, Bremen, Germany) operating in negative ion mode, as described by Dell’Acqua *et al*. (2025).

Raw LCMS data (4,758 features) were processed and curated (signal-to-noise ratio > 10; coefficient of variation in quality controls < 30%) in MS-DIAL v.4.9 with optimised parameters, yielding 3,035 features, as described by Dell’Acqua *et al*. (2025). Annotations were performed with the FragHUB database (Dablanc *et al*., 2024) and were validated wherever possible with an in-house standard database.

#### DNA extraction, amplification and sequencing

Genomic DNA was extracted from 60 mg samples of ground stem wood tissues with the DNeasy Plant Mini Kit (Qiagen, Hilden, Germany) according to the manufacturer’s instructions, with the inclusion of 12 extraction blanks (i.e. containing no sample). The DNA concentration of all samples was standardised at a maximum of 15 ng/µL after quantification on a DS-11 spectrophotometer (DeNovix, Wilmington, USA).

Fungal DNA was specifically amplified by targeting the ITS region of the fungal rRNA operon with the ITS1f/ITS2 primer pair (CTTGGTCATTTAGAGGAAGTAA/ GCTGCGTTCTTCATCGATGC). We used the amplification protocol described by Dell’Acqua *et al*. (2025). We included 30 PCR blanks (i.e. without DNA template) and 22 positive controls (11 pure DNA extracts each for *Candida oceani* and *Yamadazyma barbieri*, as described by Fournier *et al*., 2025). The quality and specificity of amplification were controlled by electrophoresis in a 2% agarose/TAE (m/v) gel to visualise the PCR products.

Bacterial DNA was specifically amplified by targeting the V5/V6 region of the 16S RNA gene with the chloroplast-excluding primer pair 799f/1115r (AACMGGATTAGATACCCKG/AGGGTTGCGCTCGTTG). Except for the primers, the amplification protocol used was identical to that for fungal DNA amplification. We included 34 positive controls (8 pure extracts of *Bacillus thuringiensis* and 26 pure extracts of *Erwinia persicina*, courtesy of Manon Chargy [SAVE]).

PCR products were sent to the Genome 706 Transcriptome Facility of Bordeaux (Cestas, France; Grants from Investissements d’Avenir, *Convention 707 attributive d’aide EquipEx Xyloforest* ANR-10-EQPX-16-01) for sequencing on an Illumina MiSeq platform (v3 chemistry, 2×300 bp). PCR product purification, the addition of multiplex identifiers and sequencing adapters, library sequencing and sequence demultiplexing (with exact index search) were performed separately for fungal and bacterial metabarcoding by the sequencing service.

#### Bioinformatics analyses

We performed bioinformatic analyses and constructed OTU tables with the FROGS pipeline implemented on a Galaxy server (https://vm-galaxy-prod.toulouse.inrae.fr/galaxy_main/; Escudié *et al*., 2018). Data preparation, clustering and chimera removal were performed as described by Dell’Acqua *et al*. (2025). A taxonomic annotation of OTUs was performed. The affiliations of each individual OTU were checked manually and, if necessary, were modified according to the following workflow. BLAST affiliation based on ITS UNITE Fungi 8.3 is considered the default reference for ITS data. BLAST affiliation based on 16S SILVA 138.1 pintail 100 is considered the default reference for 16S data. In cases of (i) disagreements or (ii) imprecise affiliations (multi-affiliation or unidentified), we performed BLAST affiliations against the NCBI (nr/nt) nucleotide collection (https://blast.ncbi.nlm.nih.gov/Blast.cgi?PROGRAM=blastn&PAGE_TYPE=BlastSearch&LINK_LOC=blasthome) or trunkdiseaseID.org (https://www.grapeipm.org/d.live/?q=td-lab-dna). NCBI was considered to be the reference of last resort.

### Data analysis

#### Statistical tools

All analyses were performed with *R* v.4.2.1. Modelling was performed through Linear Mixed Modelling (LMM) procedures implemented in the *lme4* package. All models were checked graphically for the normality of the residuals (QQ-plot) and the homogeneity of the residual variance (residuals *vs*. fitted, residuals *vs*. predictors). Otherwise, non-parametric alternatives were used or the data were transformed.

Pearson’s tests were performed and visualised with the *corrplot* package. Metabarcoding data were studied with the *phyloseq* and *vegan* packages. Metabolomic data were analysed in *Metaboanalyst* v.6.0 and *R* v.4.2.1. All tests were performed at the 5% level of significance (α = 0.05).

#### Effect of cultivar on ecophysiological and anatomical traits

We first analysed the effect of cultivar on constitutive ecophysiological traits (i.e. g_s_ and g_min_ of control plants; Supplementary Table S3) and then the effects of mean stomatal conductance, minimum leaf conductance per cultivar and their interaction on the mean incidence of esca foliar symptoms in 2024. We then investigated the effect of cultivar and stem health status on the wood anatomical traits measured in particular zones on cross-sections of xylem (i.e. specific K_th_, *F*_RP_, *F*_AP_, number of RP layers, A_flumen_ and N_P_ ; Supplementary Table S4). We performed Pearson’s correlation tests to analyse the relationship between these varietal constitutive traits and varietal mean esca susceptibility (i.e. mean esca foliar symptom incidence, at the cultivar level, over the 2017-2024 period; Fig. 1).

In addition, we tested the relation between pit density/aperture fraction and xylem vulnerability to cavitation (Ψ_50_) using the data presented in Lamarque *et al*. (2023).

#### Effect of cultivar on plant physiological response to esca

We first assessed the effects of stem health status, cultivar and their interaction on log1p- transformed occlusion PLC_th_ and P_starch_. We assessed the effect of the presence of vascular occlusions in vascular bundles bordering a given parenchyma ray on the log1p-transformed proportion of the area of the ray storing starch. On an extended subsample of esca/tiger-stripe stems from 32 cultivars, we tested the effects of esca foliar symptom severity and esca varietal susceptibility on the log1p-transformed proportion of occluded vessels. We also characterised the impact of esca expression on gas exchange by independently modelling g_s,max_ and A_max_ according to health status, cultivar and their interaction. Details on statistical procedures are provided in Supplementary Table S5. Finally, we performed Pearson’s correlation tests to assess the relationships between these varietal response-related traits and varietal mean esca susceptibility (Fig. 1).

#### Effect of esca and cultivar on the stem metabolome

Normalisation against the sample median, square-root data transformation and Pareto scaling were applied to the metabolomic data before each analysis. We performed a PERMANOVA procedure, followed by pairwise comparisons, to investigate the effect of esca varietal susceptibility cluster on the structure of the stem metabolome in control plants. We then identified differentially abundant metabolites for each pairwise comparison in Volcano plot analyses, with thresholds of a fold change > 2 and a *p*-value < 0.01. Putative contaminants (i.e. non-plant metabolites) were manually curated based on published findings and our own knowledge (Supplementary Table S2). The differentially abundant chemical features were assigned to chemical taxa with InChiKeys, using the structural ontology tool ClassyFire (Djoumbou Feunang *et al*., 2016) and the interface ClassyFire Batch (https://cfb.fiehnlab.ucdavis.edu/). We compared the enrichment in specific structural ontologies graphically for each pairwise comparison.

A similar procedure was performed separately for each esca varietal susceptibility cluster, to evaluate the effect of stem health status on the stem microbiome (Fig. 1). A similar procedure was performed on esca/tiger-stripe stems to evaluate the effect of esca symptom severity on the stem microbiome.

We also investigated the effect of stem health status on metabolomic variance by testing for the homogeneity of multivariate dispersions (PERMDISP) through calculations of the mean distance to centroids based on Euclidean distances and 999 permutations. Pairwise post-hoc comparisons were performed if the global test yielded significant results.

#### Effect of esca and cultivar on stem microbial communities

All analyses were performed separately for fungal and bacterial data. Before the statistical analysis of the metabarcoding data, we decontaminated the dataset (removing putative contaminant OTUs and highly contaminated samples) with the tools available in the metabaR package (Zinger *et al*., 2021).

We evaluated the effect of esca varietal susceptibility cluster on the constitutive diversity and structure of stem microbial communities in control plants (Fig. 1). We compared the Shannon diversity index between clusters. We investigated effects on microbial community structure by performing a PERMANOVA on CLR-transformed data with the addition of pseudocounts of one. We checked the multivariate homogeneity of dispersions before performing pairwise comparisons. Similar processes (i.e. Shannon index comparisons and PERMANOVA) were performed (i) to evaluate the effects of stem health status, esca varietal susceptibility cluster and their interaction on the diversity and structure of stem microbial communities in all plants (Fig. 1); and (ii) to evaluate the effect of esca symptom severity on the diversity and structure of stem microbial communities in esca/tiger-stripe stems.

## Results

### Weakly susceptible cultivars exhibit low stomatal conductance

We observed a significant cultivar effect on g_s_ (*P <* 10^-16^), which ranged from 97.6 ± 7.6 mmol m^-2^ s^-1^ to 249.4 ± 19.9 mmol m^-2^ s^-1^ (Supplementary Fig. S1A; Supplementary Table S3). At cultivar level, g_s_ was not significantly correlated with the incidence of esca foliar symptoms in the next summer season (*r =* 0.25, *P =* 0.09; Fig. 3A). However, interestingly, none of the eight cultivars with the lowest g_s_ values had an incidence of esca foliar symptoms above 30% in 2024 (Fig. 3A; Supplementary Fig. S1A). The overall correlation of g_s_ with varietal mean esca susceptibility over the period 2017-2024, as indicated by the colour gradient in Fig. 3A, was of borderline significance (r = 0.28, *P* = 0.06).

**Figure 3.**
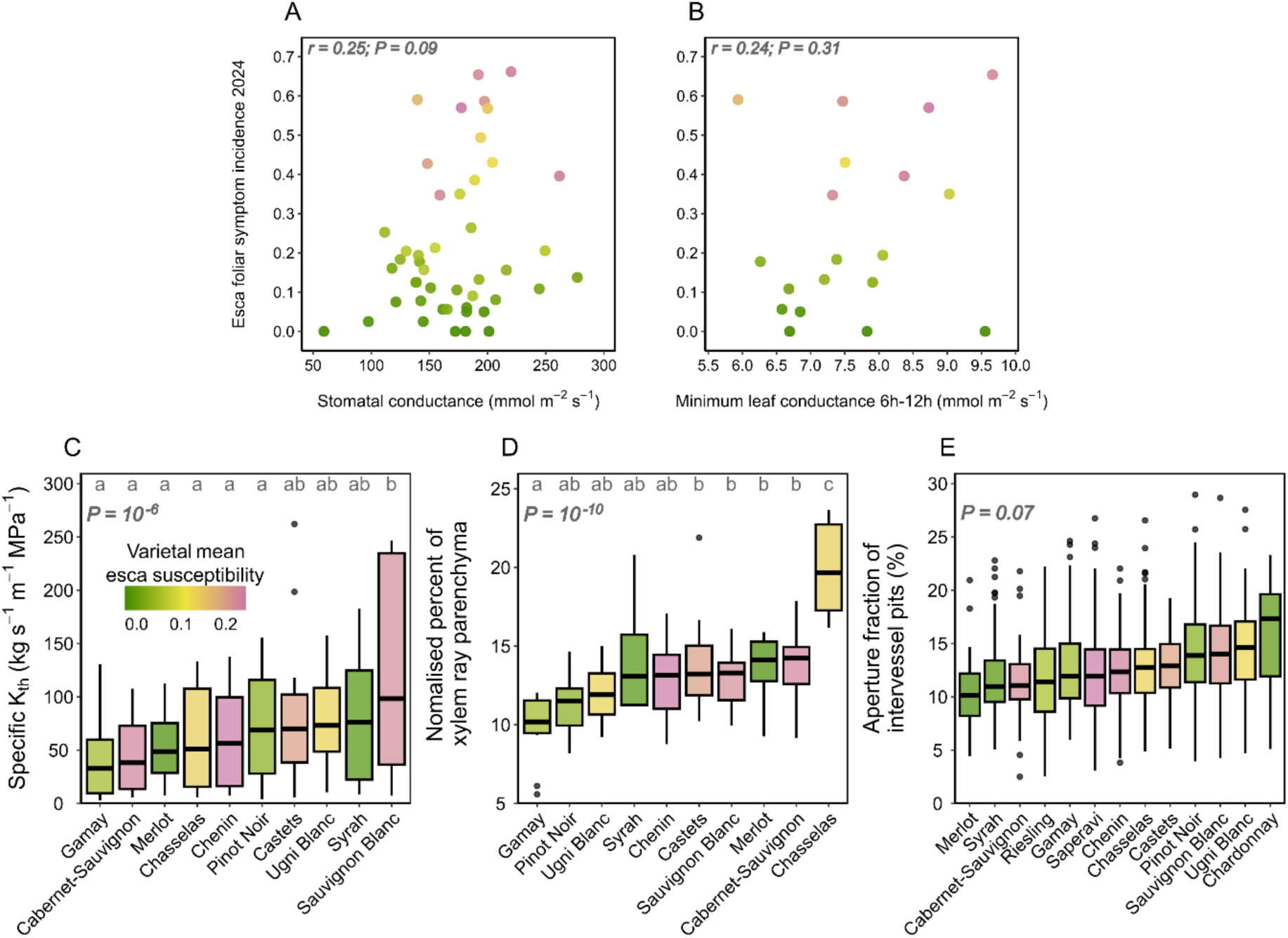
Varietal relationships between leaf gas exchange, stem xylem anatomy and varietal esca susceptibility. (A) Correlation between varietal mean stomatal conductance measured in June 2024 and esca foliar symptom incidence in the summer 2024 on 46 cultivars (*n* = 1,840 plants) (B) Correlation between varietal mean minimum leaf conductance measured in May 2024 on detached leaves and esca foliar symptom incidence in the summer 2024 on 19 cultivars (*n* = 114 leaves). The relationship was fitted to the data for 18 cultivars (after excluding Cabernet Franc): *cloglog (Esca foliar symptom incidence 2024) = -6.19 + 1.54 × g_min_ (*P < *10^-16^)*. Each dot corresponds to the value obtained for a specific cultivar. (C) Boxplots representing mean specific theoretical hydraulic conductivity per zone (i.e. three vascular bundles) across 10 cultivars (*n* = 60 stems) (D) Boxplots representing the normalised percentage of stem xylem ray parenchyma for 10 cultivars (*n* = 60 stems) (E) Boxplots representing the aperture fraction of intervessel pits for 13 cultivars (*n* = 42 stems). Boxplots display the median and interquartile range, with whiskers extending to the minimum and maximum values, excluding outliers, which are shown as individual black points. Colours indicate varietal mean esca susceptibility (mean foliar symptom incidence from 2017 to 2024). Correlation coefficients and associated *p-*values were obtained in Pearson’s tests. *P*- values correspond to cultivar effects in linear mixed models. The letters correspond to significance groups in Tukey tests with an alpha risk of 5%.

We also observed a significant variation of g_min_ between cultivars (*P =* 10^-7^): g_min_ ranging from 5.9 ± 0.32 mmol m^-2^ s^-1^ to 9.7 ± 0.53 -mmol m^-2^ s^-1^ (Supplementary Fig. S1B; Supplementary Table S3). At cultivar level, g_min_ was not significantly correlated with the incidence of esca foliar symptoms in the next summer season (*r =* 0.24, *P =* 0.31; Fig. 3B). In addition, g_min_ was not significantly correlated with varietal mean esca susceptibility over the period 2017-2024 (*r =* 0.28, *P =* 0.25; Fig. 3B). However, there was a trend towards a positive correlation for most cultivars, with the exception of two extreme phenotypes: Cabernet Franc (low g_min_ and high esca susceptibility) and Xinomavro (high g_min_ and low esca susceptibility).

Finally, g_s_ and g_min_ were not significantly correlated at varietal level (*r =* 0.25, *P =* 0.29).

### The genetic diversity of wood anatomical traits is not correlated with varietal esca susceptibility

Specific K_th_ was significantly influenced by cultivar but not by stem health status (*P =* 10^-6^, *P =* 0.58, respectively). Specific K_th_ ranged from 41.7 ± 8.0 kg s^-1^ m^-1^ MPa^-1^ to 120.5 ± 20.0 kg s^-1^ m^-1^ MPa^-1^ (Fig. 3C). Specific K_th_ was not significantly correlated with varietal mean esca susceptibility (*r =* 0.27, *P =* 0.45; Fig. 3C). *F*_RP_ was significantly influenced by cultivar but not by stem health status (*P =* 10^-10^, *P =* 0.61, respectively). *F*_RP_ ranged from 9.9 ± 0.4 % to 20.4 ± 0.8 % (Fig. 3D). *F*_RP_ was not significantly correlated with varietal mean esca susceptibility (*r =* 0.14, *P =* 0.69; Fig. 3D). Cultivar also had a significant effect on the number of RP layers *(P =* 10^-6^; Supplementary Fig. S2A) and A_flumen_ (*P =* 10^-4^; Supplementary Fig. S2B). Neither the number of RP layers nor A_flumen_ was significantly correlated with varietal mean esca susceptibility (*r =* 0.09, *P =* 0.81 and *r =* 0.27, *P =* 0.45 Supplementary Fig. S2A and B). In esca/tiger-stripe stems, the severity of esca foliar symptoms did not significantly affect specific K_th_, *F*_RP_, number of RP layers or A_flumen_ (*P =* 0.06, *P =* 0.99, *P =* 0.65 and *P =* 0.77, respectively).

Cultivar had an effect of borderline significance on *F*_AP_ (*P =* 0.07, *P =* 0.06, respectively). *F*_AP_ ranged from 10.3 ± 0.6 % to 15.9 ± 0.4 % (Fig. 3E). *F*_AP_ was not significantly correlated with varietal mean esca susceptibility (*r =* -0.05, *P =* 0.88; Fig. 3E). N_P_ was not significantly influenced by cultivar (*P =* 0.41) and not significantly correlated with varietal mean esca susceptibility (*r =* -0.17, *P =* 0.59; Supplementary Fig. S2C). Detailed statistics for wood anatomical traits are provided in Supplementary Table S4.

### Co-occurrence of impairments of hydraulic conductivity, starch storage and gas exchange in symptomatic stems of grapevine cultivars

Occlusion PLC_th_ was significantly influenced by stem health status, but not by cultivar or their interaction (*P =* 10^-9^, *P =* 0.71, *P =* 0.73, respectively). There was almost no loss of theoretical hydraulic conductivity due to occlusion in control stems (0.6 ± 0.2 %; Fig. 4A), whereas a mean loss of 22.1 ± 2.9% was found in esca/tiger-stripe stems (Fig. 4B). For these stems, there was no significant difference between the 10 cultivars, although occlusion PLC_th_ ranged from 11.7 ± 8.3% to 38.8 ± 11.7 % (Fig. 4B). In addition, occlusion PLC_th_ in esca/tiger-stripe stems was not significantly correlated with varietal mean esca susceptibility (*r =* 0.02, *P =* 0.68; Fig. 4B) and not significantly influenced by the severity of esca foliar symptoms (*P =* 0.81).

**Figure 4.**
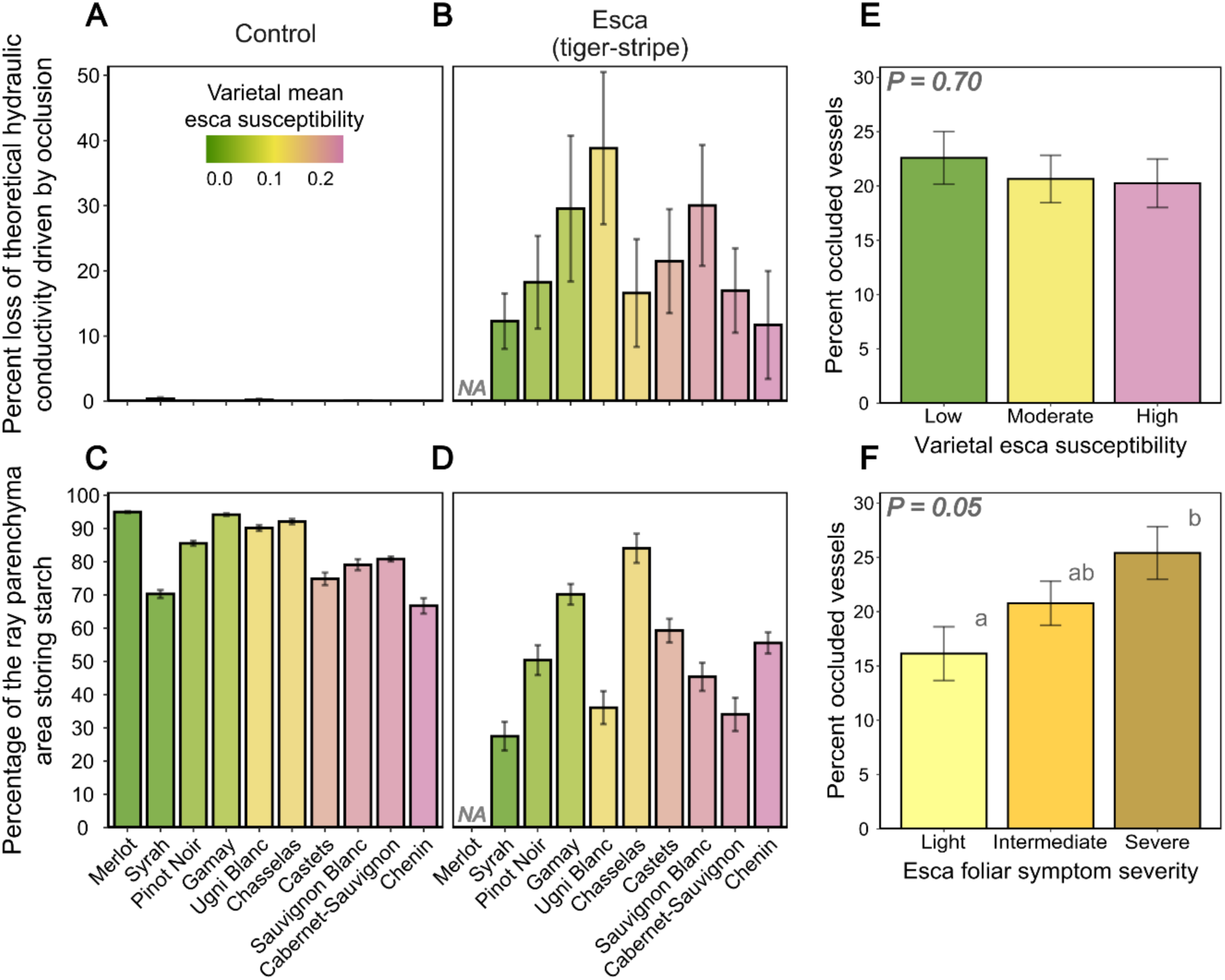
Vascular functional responses to esca across differentially susceptible cultivars. (A, B) Percent loss of theoretical hydraulic conductivity due to vascular occlusions (mean ± SEM) in 10 cultivars for (A) control stems (*n* = 35) and (B) esca/tiger-stripe stems (*n* = 25). (C, D) Percentage of the ray parenchyma area storing starch (mean ± SEM) in 10 cultivars for (C) control stems (*n* = 35) and (D) esca/tiger-stripe stems (*n* = 25). (E, F) Percentage of occluded vessels in esca/tiger-stripe stems for (E) varietal esca susceptibility classes (*n* = 32 cultivars, *n* = 81 stems) and (F) esca foliar symptom severity levels. Barplots are colour-coded and cultivars are ordered by varietal mean esca susceptibility (mean foliar symptom incidence between 2017 and 2024) or esca foliar symptom severity (F). *P*-values correspond to effects in linear mixed models. The letters correspond to significance groups in Tukey tests with an alpha risk of 5%.

We extended the range of cultivars studied for the occurrence of esca-associated vascular occlusions by determining the percentage of occluded vessels in 81 esca/tiger-stripe stems from 32 cultivars. The percent occluded vessels in esca/tiger-stripe stems was not significantly influenced by varietal mean esca susceptibility class (*P =* 0.70; Fig. 4E). However, there was considerable variability within cultivars, making it impossible to identify trends. The percent occluded vessels was marginally influenced by the severity of esca foliar symptoms (*P =* 0.05), with more occlusions in stems with severe symptoms than in stems with light symptoms (Fig. 4F).

P_starch_ was significantly affected by stem health status, but not by cultivar or the interaction of these two factors (*P =* 10^-6^, *P =* 0.13, *P =* 0.45, respectively). It was lower in esca/tiger-stripe stems than in control stems (51.3 ± 1.5 % *vs*. 83.7 ± 0.5 %; Fig. 4C and D). P_starch_ in control stems did not differ significantly between the 10 cultivars (Fig. 4C). It was also not significantly correlated with varietal mean esca susceptibility, despite the observation of a negative trend (*r =* -0.44, *P =* 0.20; Fig. 4C). P_starch_ in esca/tiger-stripe stems did not differ significantly between the 10 cultivars (Fig. 4D). It was not significantly correlated with varietal mean esca susceptibility (*r =* 0.07, *P =* 0.85; Fig. 4C). P_starch_ in esca/tiger-stripe stems was not significantly influenced by the severity of esca foliar symptoms (*P =* 0.21).

As (i) the variability of P_starch_ was very high in esca/tiger-stripe stems (RSD = 67.5% *vs.* 16.1% in control stems; Table 1), and (ii) this trait was strongly correlated with occlusion PLC_th_ in stems (*r =* -0.71, *P =* 10^-10^), we tested the effect of the occurrence of vascular occlusions in vascular bundles bordering a given parenchyma ray on the percentage of the ray area storing starch. P_starch_ in esca/tiger-stripe stems was significantly influenced by the occurrence of vascular occlusions (*P =* 10^-10^). It was significantly lower when occlusions were present in a single adjacent bundle, and even lower when there were occlusions in both adjacent bundles (Fig. 2B, C, E and F, Supplementary Fig. S3). There was a global trend, across all cultivars, towards a decrease in P_starch_ in the presence of occlusions, whereas the difference between one and two occluded bundles was less consistent between cultivars (Supplementary Fig. S4).

We hypothesised that the decrease in starch storage in esca/tiger-stripe stems might be related to an impairment of carbon assimilation in symptomatic plants (Bortolami *et al*., 2021b). We therefore measured gas exchange before and during the expression of esca leaf symptoms in plants of different subsequent health statuses (i.e. control *vs*. esca/tiger-stripe stems) in a subset of five cultivars. Before esca expression, g_s,max_ was not significantly influenced by upcoming health status (*P =* 0.99; Supplementary Fig. S5A). A_max_ was not significantly influenced by health status (*P =* 0.76; Supplementary Fig. S5B). During esca expression, g_s,max_ was significantly influenced by cultivar and health status (*P =* 10^-5^, *P =* 10^-6^, respectively). A_max_ was significantly influenced by health status, but not by cultivar (*P =* 10^-6^, *P =* 0.12, respectively). Both g_s,max_ and A_max_ were significantly lower on green leaves from esca/tiger-stripe stems (Supplementary Fig. S5C and D). Detailed statistics for stem physiological responses to esca are provided in Supplementary Table S5.

### Neither esca expression nor cultivar susceptibility alters the structure and diversity of stem microbial communities

In total, 2,670,894 fungal reads were obtained, distributed across 242 OTUs. There was a mean numbe r of 30,700 ± 73 fungal reads per sample (minimum: 9,444; maximum: 53,800). Three fungal phyla were retrieved across the 10 cultivars: Ascomycota (84% of fungal reads), Basidiomycota (15%), and Chytridiomycota (<1%). The three most abundant fungal genera were *Aureobasidium* sp. (34.8% of fungal reads), *Cladosporium* sp. (25.5%), and *Alternaria* sp. (14.5%; Supplementary Fig. S6). In total, 292,439 bacterial reads were obtained, distributed across 512 OTUs. There was a mean number of 3,361 ± 11 bacterial reads per sample (minimum: 1,343; maximum: 7,738). We identified 17 bacterial phyla, predominantly Proteobacteria (63% of bacterial reads), Firmicutes (13%), and Actinobacteriota (12%). The three most abundant bacterial genera were *Ralstonia* sp. (18.5% of bacterial reads), *Stenotrophomonas* sp. (17.4%), and *Acinetobacter* sp. (7.7%; Supplementary Fig. S7). Rarefaction curves approached saturation for most samples, indicating sufficient sampling of dominant taxa (Supplementary Fig. S8). Decontaminated OTU tables are provided in Supplementary Table S6 (ITS) and Supplementary Table S7 (16S).

We first compared the microbial communities present in control stems across varietal susceptibility classes, with cultivar as a covariate. Varietal esca susceptibility did not significantly influence fungal or bacterial Shannon diversity indexes (Supplementary Fig. S9A and B). Varietal esca susceptibility did not significantly modify fungal or bacterial community structure (7*P* = 0.20; *P* = 0.26; Supplementary Fig. S9C and D), whereas cultivar did have an effect (28.1% explained variance, *P* = 0.03; 28.4% explained variance, *P* = 0.01).

Shannon diversity index was not significantly influenced by stem health status and varietal esca susceptibility for either fungal (*P* = 0.87, *P =* 0.94) or bacterial communities; *P* = 0.11, *P =* 0.57 ; Fig. 5A and C). Neither stem health status nor varietal esca susceptibility significantly affected fungal (*P* = 0.33; *P* = 0.34; Fig. 5B) or bacterial (*P* = 0.18; *P* = 0.14; Fig. 5D) community structure, whereas cultivar did have an effect (9.6% explained variance, *P* = 0.03 for fungi; 9.3% explained variance, *P* = 0.02).

**Figure 5.**
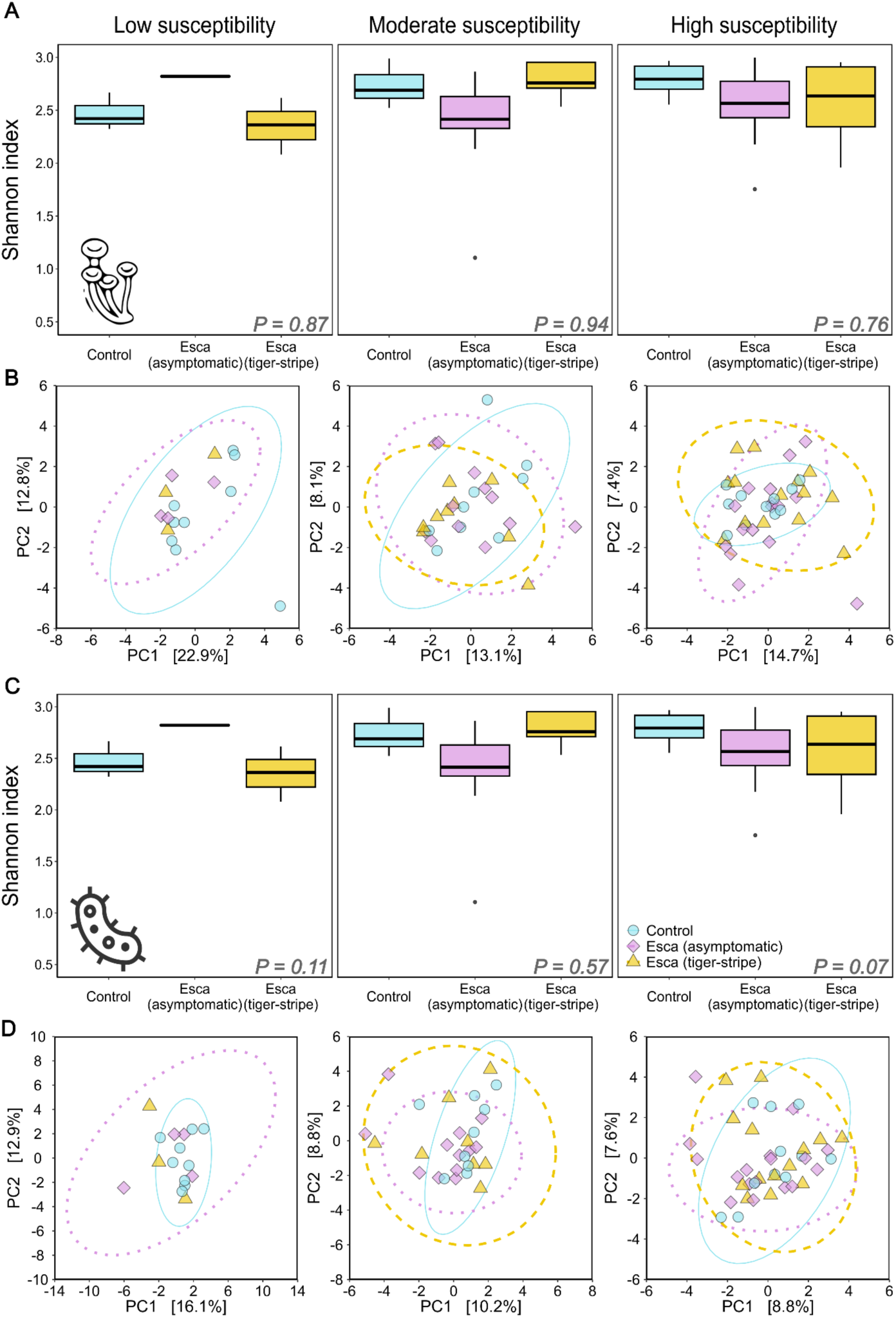
Effects of esca expression on stem microbial communities and relationship to varietal esca susceptibility. (A, **C)** Effect of esca expression and esca varietal susceptibility class on the Shannon index associated with (A) fungal and (C) bacterial communities, for weakly susceptible (*n* = 3 cultivars, *n* = 16 stems), moderately susceptible (*n* = 3 cultivars, *n* = 30 stems), and highly susceptible (*n* = 4 cultivars, *n* = 41 stems) cultivars. Boxplots displaying the median and interquartile range, with whiskers extending to the minimum and maximum values, excluding outliers, which are shown as individual black points. **(B, D)** PCA summarising the effects of esca expression and esca varietal susceptibility class on the structure of (B) fungal and (D) bacterial communities calculated with CLR-transformed data, for weakly susceptible, moderately susceptible, and highly susceptible cultivars. Ellipses correspond to the 95% confidence interval for each group. The colour and shape of the points indicate stem health status. *P*-values correspond to esca effects in ANOVA analyses on Shannon index.

Shannon diversity index was not significantly modified by esca foliar symptom severity in esca/tiger-stripe stems (Supplementary Fig. S10A and C). Foliar symptom severity did not significantly alter the structure of microbial communities (Supplementary Fig. S10B and D). Detailed statistics for stem microbial communities are provided in Supplementary Table S8.

### Genotypic differences in the constitutive stem metabolome are not clearly related to esca susceptibility

Untargeted metabolomics yielded 3,035 curated metabolic features (see Materials and Methods), which were used for the metabolome analysis. We first compared the metabolome of control stems between varietal susceptibility classes. Varietal esca susceptibility significantly modified the structure of the overall metabolome (15.4% explained variance, *P* = 0.001; Supplementary Fig. S11A). Moderately susceptible cultivars displayed enrichment in a larger number of compounds than cultivars of lower susceptibility (21 *vs.* 8 features; Supplementary Fig. S11B and C). A clear enrichment in “phenylpropanoids and polyketides” (especially flavonoids) was observed in moderately susceptible cultivars relative to weakly susceptible cultivars (9 *vs.* 1 features), whereas no clear enrichment was observed in comparisons of weakly and highly susceptible cultivars (7 features each), or of moderately and highly susceptible cultivars (15 features each; Supplementary Fig. S11E, F and G). Flavonoids were overabundant in moderately susceptible cultivars and other phenylpropanoids (e.g. 3- hydroxyphysodic acid, a phenolic glycoside and a kavalactone) were overabundant in weakly susceptible cultivars. A very slight enrichment in “terpenes” was observed in moderately and highly susceptible cultivars (Supplementary Fig. S11F and G). Other secondary metabolites (i.e. neochlorogenic acid and a benzenesulfonate ester) were found specifically among the top-ranked markers for high susceptibility (log*_2_*FC > 8).

### Induced stem metabolic response to esca involves a strong accumulation of specialised metabolites and lipids in susceptible cultivars

We found a significant effect of esca expression on the structure of the metabolome for all varietal susceptibility classes. The percentage of the variance explained by stem health status decreased with increasing cultivar susceptibility (23.5% explained variance in cultivars with low susceptibility to esca, *P* = 0.008; 12.1% explained variance in moderately susceptible cultivars, *P* = 0.02; 8.6% explained variance in highly susceptible cultivars, *P* = 0.01; Fig. 6A, B, and C; Supplementary Fig. S12A; Supplementary. Fig S13A). We found that health status affected metabolite distribution in highly (*P* = 0.01) and moderately susceptible cultivars (*P* = 0.02), but not in weakly susceptible cultivars (*P* = 0.17) (Fig. 6A, B and C).

**Figure 6.**
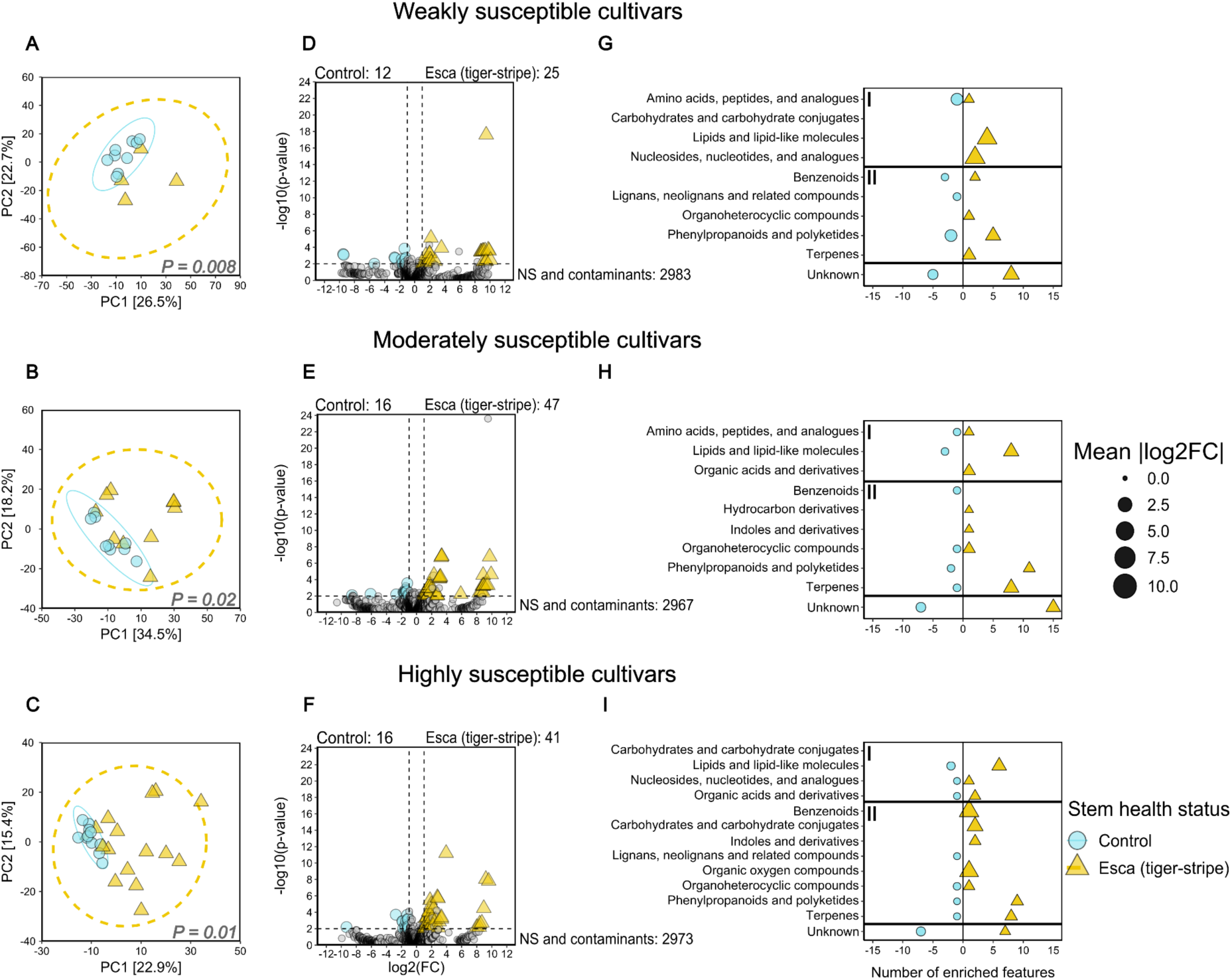
Effects of esca expression on the stem metabolome and relationship to esca varietal susceptibility. Panels are grouped by esca susceptibility: (A, D, G) weakly susceptible cultivars (*n* = 3 cultivars, *n* = 13 stems), (B, E, H) moderately susceptible cultivars (*n* = 3 cultivars, *n* = 19 stems), and (C, F, I) highly susceptible cultivars (*n* = 4 cultivars, *n* = 26 stems). **(A, B, C)** PCA summarising the effects of esca expression on the structure of stem metabolome performed on normalised, transformed and scaled data. Ellipses correspond to the 95% confidence interval for each group. Dots and ellipses are coloured and shaped according to the stem health status. **(D, F, E)** Volcano plots on all features. Cut-off values for significance were set at |log_2_FC| > 2 and *P* < 0.01. Non-significant (NS) and putatively contaminant features are shown in grey. Features for which enrichment was observed in control stems are shown as blue circles, and features for which enrichment was observed in esca/tiger-stripe stems are shown as yellow triangles. **(G, H, I)** Class assignment of differentially abundant features selected in Volcano analyses (cut-off values set at |log_2_FC| > 2 and *P* < 0.01). Class assignment was performed with Classyfire and published findings. Classes for which enrichment was observed in control stems are shown as blue circles, and classes for which enrichment was observed in esca/tiger-stripe stems are shown as yellow triangles. Features that could not be assigned to any known compound or metabolic class are classified as “Unknown”. Metabolic classes were classified as belonging to primary (I, above) or secondary (II, below) metabolism.

The number of metabolites displaying enrichment in response to esca was greater in moderately and highly susceptible cultivars (47 and 41 differentially abundant features, respectively) than in weakly susceptible cultivars (25 differentially abundant features; Fig. 6D, E and F). An enrichment in “lipids and lipid-like molecules” and in “phenylpropanoids and polyketides” was clearly observed in esca/tiger-stripe stems relative to control stems, for all cultivars (Fig. 6G, H and I). However, a greater enrichment in “phenylpropanoids and polyketides” was observed in moderately and highly susceptible cultivars (11 and 9 features, respectively) than in weakly susceptible cultivars (5 features). A clear enrichment in “terpenes” was observed in moderately and highly susceptible cultivars (8 features each), but not in weakly susceptible cultivars (1 feature) (Fig. 6H and I). Primary metabolites (lipids and pyrimidine nucleosides) were specifically represented among the top-ranked markers of esca (log_2_FC > 8) in weakly susceptible cultivars. Terpenes and lipids were specifically represented among the top-ranked markers of esca in moderately susceptible cultivars. Interestingly, specialised glycosylated secondary metabolites (i.e. terpene glycoside, phenolic glycosides and a derivative of alpha-D-glucopyranoside) were specifically represented among the top-ranked markers of esca in highly susceptible cultivars.

There were no clear differences between control stems and esca/asymptomatic stems (Supplementary Fig. S12). The differences between esca/asymptomatic and esca/tiger-stripe stems were similar to those between control and tiger-stripe stems (Supplementary Fig. S13).

Finally, esca foliar symptom severity did not significantly affect metabolome structure in esca/tiger-stripe stems (12.0% explained variance, *P* = 0.13; Supplementary Fig. S14A). However, enrichment was observed for a larger number of compounds in stems displaying severe symptoms than in stems with intermediate symptoms (28 *vs.* 15 features). This was the case, particularly, for “phenylpropanoids and polyketides” (13 *vs.* 2 features; Supplementary Fig. S9C, D, F and G). No enrichment in any particular metabolic class was observed in stems displaying severe symptoms relative to those with mild symptoms (18 *vs.* 10 features; Supplementary Fig. S14E and G), or in stems with intermediate symptoms relative to those with mild symptoms (5 *vs.* 5 features; Supplementary Fig. S14E and F).

## Discussion

In this study, we showed that intraspecific variation in susceptibility to esca, a complex vascular disease, is primarily associated with plant traits rather than wood microbial communities. Across 46 grapevine cultivars grown in a common garden, conservative water-use strategies were associated with lower esca susceptibility. Esca induced a conserved syndrome of vessel occlusion likely leading to hydraulic dysfunction and carbon depletion across cultivars, but divergent metabolic strategies: highly susceptible genotypes displayed stronger induction of some defence metabolic pathways, potentially contributing to foliar symptom expression.

### Conservative water-use strategies are associated with weak susceptibility to esca

Hydraulic functioning is increasingly recognised as a key determinant of plant responses to vascular diseases and woody plant decline (Yadeta & Thomma, 2013; Torres-Ruiz *et al*., 2024). In line with our initial hypothesis, most of the grapevine cultivars with foliar water-saving strategies (i.e. low stomatal conductance and/or a low minimum conductance) displayed low susceptibility to esca. This result is consistent with the total inhibition of esca foliar symptoms under drought, during which stomatal conductance and overall transpiration of the canopy are reduced (Bortolami *et al*., 2021b). A similar relationship has also been observed in *Eutypa* dieback in grapevine, to which cultivars tightly regulating gas exchanges have been shown to be more tolerant (Sinclair *et al*., 2025). Together, efficient water conservation and reduced vegetative growth (Gastou *et al*., 2024) may create physiological conditions less favourable to esca expression. Lower transpiration rates may limit the production of phytotoxic metabolites and their translocation from the trunk (where wood decay processes occur) to the leaves *via* the vascular system. It could also limit the activity and aggressiveness of some pathogenic fungal species (Chambard *et al*., 2026).However, this conservative strategy was not shared by all esca- tolerant varieties, as some of the genotypes studied combined low esca susceptibility and moderate-to-weak water-saving strategies (e.g. Xinomavro, Sangiovese). The use of genotypes combining strong water-saving and low esca susceptibility would constitute a win–win strategy for efforts to plant and breed multi-tolerant grapevine genotypes, especially as genotypes exhibiting a trait syndrome with lower stomatal conductance could also be more tolerant to drought (Dayer *et al*., 2022).

Contrary to our hypothesis, the relationship between water-use strategy and esca susceptibility was not underpinned by constitutive differences in xylem anatomy. We can therefore reject the hypothesis that anatomical characteristics limiting sap flow (leading to low theoretical hydraulic conductivity) and xylem connectivity (e.g., a lower proportion of ray parenchyma and a lower aperture fraction of intervessel pits) may decrease the transport of toxic metabolites and thus inhibit the expression of foliar esca symptoms. This result contrasts with the role of xylem anatomy in drought tolerance (Zhang *et al*., 2021; Liu *et al*., 2024) and in host susceptibility to the grapevine trunk pathogen *Phaeomoniella chlamydospora* (Pouzoulet *et al*., 2017, 2020) or other vascular pathogens (e.g. *Xylella fastidiosa* and *Ophiostoma novo-ulmi*; Chatelet *et al*., 2011; Venturas *et al*., 2013). As pit density/aperture fraction was not a relevant predictor of neither esca susceptibility nor Ψ_50_ (xylem vulnerability to cavitation; Supplementary Fig. S15), the characterisation of pit membrane thickness and ultrastructure (Delzon *et al*., 2010; Li *et al*., 2016) might now be useful for determining their contributions to both esca and drought vulnerability in grapevine.

### A conserved syndrome of physiological responses to esca across genotypes, despite no change in microbial community

Plant vascular diseases have severe physiological consequences that differ between genotypes according to their susceptibility (Silva *et al*., 2020; Fanton *et al*., 2022). Contrary to our second hypothesis, we highlighted a conserved syndrome of esca-induced physiological responses across cultivars. Occlusions (i.e. non-gaseous embolism) accumulated in xylem vessels, impeding theoretical stem hydraulic conductivity in all cultivars (as shown in leaves in Bortolami *et al*., 2023). To maintain their productivity in subsequent years, grapevine partially compensates for these hydraulic dysfunctions by producing new functional vessels later in the season (Dell’Acqua *et al*., 2024). Future comparisons of these mechanisms across genotypes would allow testing whether there is a trade-off between short-term response and long-term resilience to esca disease.

We also demonstrated the existence of a positive relationship between the percentage of vessels occluded in stems and the severity of leaf symptoms, as reported in other pathosystems including *Verticillium* wilt in olive tree (Trapero *et al*., 2018) and laurel wilt in avocado (Inch *et al*., 2012), providing evidence for a direct relationship between hydraulic dysfunction and symptom expression (Bortolami *et al*., 2019, 2023). Esca expression was also associated with lower levels of starch storage in the stem xylem ray parenchyma, co-occurring in the same stem sectors with vascular occlusions as previously reported for Pierce’s disease (Ingel *et al*., 2021). However, the mechanisms underlying esca symptom severity or incidence are likely different as we found here no direct link between the cultivar susceptibility to esca and the level of vascular occlusion in response to esca, in line with the limited correlation between esca foliar symptom incidence and averaged severity across cultivars (Gastou *et al*., 2024). In addition, we found no evidence for greater starch accumulation or depletion in stems of highly susceptible genotypes expressing esca symptoms, whereas both these trends have been described in other diseases (Silva *et al*., 2020; Fanton *et al*., 2022). As a putative cause of starch depletion, we observed a decrease in leaf gas exchanges only after the onset of esca leaf symptoms, slowing the build-up of carbon reserves(Zufferey *et al*., 2012; Tixier *et al*., 2019). Starch may be remobilised directly *via* the shikimate pathway to produce pectocellulosic constituents of gels and gums, or the secondary metabolites, assuming a defence/storage trade-off (Viiri *et al*., 2001; Thalmann & Santelia, 2017; Ingel *et al*., 2021). Alternatively, changes to phloem formation, integrity and functionality (e.g. obstruction of sieve plates, repression of cambial activity, starch hydrolysis) in the most affected area of the stem may also occur (Partelli-Feltrin *et al*., 2023; Prats *et al*., 2023; Moret *et al*., 2024).

On the other hand, we found that esca expression had no effect on stem microbial communities. This finding is consistent with results previously obtained *in natura* for stem fungal communities (Del Frari *et al*., 2019), although a slight decrease in Ascomycota diversity was previously reported in the young organs of potted mature vines (Gastou *et al*., 2026). However, these results are also surprising, because vascular diseases induce major physiological changes, altering the conditions in which microorganisms must survive. In other complex pathosystems, such as olive knot disease, and in response to drought, the diversity and structure of microbial communities are more plastic. This plasticity is genotype-dependent, driving genetic susceptibility to the stress concerned (Gomes *et al*., 2019; Brunschwig *et al*. 2024). The role of microbial communities in driving intraspecific varietal susceptibility therefore depends on the plant species-stress combination, and host physiology seems to make a greater contribution than microbial communities to esca.

### Contrasting defence metabolism may contribute to differences in disease expression

Contrary to our expectations, constitutive stem metabolomes did not differ consistently among cultivars with contrasting susceptibility, suggesting that basal metabolic composition alone does not explain intraspecific variation in esca susceptibility. This contrasts with recent observations in the two grapevine cultivars inoculated with *Eutypa lata* (Sinclair *et al*., 2025) and highlights the complexity of esca pathogenesis.

We then investigated whether the differences between genotypes remained similar for symptomatic stems. Esca expression did not considerably alter the overall metabolic structure of the stem, instead inducing, in all genotypes, a clear biochemical deregulation: increased variability of the metabolic structure, overexpression of lipids, terpenes, and phenylpropanoids. This response was restricted to symptomatic stems, confirming that many esca-induced disorders (occlusions, impaired gas exchange, metabolic responses) are localised along specific sap flows (Bortolami *et al*., 2021a, b), whereas others, such as DNA methylation, are systemic (Berger *et al*., 2026). The accumulation of defence-related compounds is a classical response to esca, from perennial wood to leaves (Fontaine *et al*., 2016; Berger *et al*., 2026; Chambard *et al*., 2026).

Contrary to our third hypothesis, a greater accumulation of secondary metabolites in esca-symptomatic stems was observed in moderately and highly susceptible genotypes. The biological origin of these metabolites remains unclear. They may include defence-related compounds, low-molecular weight fungal compounds (e.g. toxins, chelators), and by-products of wood degradation. Strong accumulation of such by-products in highly susceptible cultivars is probable, due to the large proportion of necrotic wood in these plants (Gastou *et al*., 2025): this could ultimately lead to an increase in the frequency of esca leaf symptoms in the long term.

The opposite result was previously reported for esca-symptomatic vines, as well as in other pathosystems, with a greater overaccumulation of antioxidant proteins, bioactive stilbenoids, fatty acids or phospholipids in weakly susceptible genotypes (Lemaitre-Guillier *et al*., 2020; Khattab *et al*., 2021; Das *et al*., 2025).

A non-exclusive explanation is that susceptible cultivars’ defence strategy differs not only in its magnitude but also in its nature. Some stress-response pathways would therefore be overexpressed in an uncontrolled manner as a response to esca in susceptible genotypes (potentially at the expense of lignin biosynthesis; Negro *et al*., 2020), either directly in the stem (i.e. distal to the infection area) or upstream (e.g. in perennial organs), and may act as a signal for frequent and severe esca symptom expression. In the same vein, a link between anthocyanin accumulation and symptom expression in susceptible cultivars was previously suggested in the *flavescence dorée* pathosystem (Margaria *et al*., 2014). Several glycosylated compounds were overexpressed in the most susceptible cultivars, as previously reported in wild grapevine inoculated artificially with the pathogenic fungus *Neofusicoccum parvum* (Khattab *et al*., 2021). The accumulation of glycosylated metabolites may reflect a shift in primary metabolism towards defence, an inability to efficiently produce bioactive phenylpropanoids, or the detoxification of overabundant phytotoxic compounds (Velasco *et al*., 2007; Le Roy *et al*., 2016).

## Conclusion

Overall, this integrative study provides new insights into the functional determinants of intraspecific susceptibility to complex vascular diseases in perennial plants. Our results indicate that disease susceptibility is primarily associated with coordinated variations in plant functional traits rather than wood microbial communities, highlighting the predominant role of host physiological strategies in shaping disease outcomes.

Building on these findings, we propose a hypothetical functional framework in which contrasting functional physiological strategies across cultivars govern disease expression by influencing the production, transport and accumulation of defence-related metabolites. This potentially reflects trade-offs between resource acquisition, water conservation and defence responses.

Testing this framework within and across perennial plant species will require multiscale studies of coordinated variations in plant physiology, metabolism, microbial interactions and environmental conditions across spatial and temporal dimensions. Such trait-based approaches may also help identify physiological targets for breeding more resilient genotypes.

## Acknowledgements

We thank Agnes Destrac Irvine and Cornelis van Leeuwen for providing access to the VitAdapt experimental vineyard and the Experimental Viticultural Unit of Bordeaux 1442, INRAE, F- 33883 Villenave d’Ornon for vineyard management. We also thank Mathéo Pinol Daubisse and the teams from SAVE, BIOGECO and EGFV units for help in field measurements. Minimum conductance measurements were performed at the PHENOBOIS platform (Pessac, France) and DNA sequencing at the Genome Transcriptome facility of Bordeaux (PGTB, Cestas, France). We thank Alexandre Chataigner, Linda Stammitti-Bert, Ana Clara Fanton, Ninon Dell’Acqua and Marie Chambard for helpful discussions and advices on molecular biology analyses, wood phenotyping, metabarcoding or metabolomics.

Pierre Gastou received a PhD grant from the French *Ministère de l’Enseignement Supérieur et de la Recherche*. This study received financial support from Château-Figeac (Saint-Emilion), the French government in the framework of the IdEX Bordeaux University “Investments for the Future” programme / GPR Bordeaux Plant Sciences, and the French National Research Agency (ANR) in the framework of the “Investments for the Future Programme”, within the Cluster of Excellence COTE (ANR-10-LABX-45).

## Competing interests

The authors have no competing interests to declare.

## Author contributions

C.E.L.D. and P.G. designed the experiment. C.E.L.D. supervised the project and secured the funds. L.A., P.G. and A.M. performed leaf gas exchange measurements. L.A. and P.G. performed leaf sampling. L.A performed minimum leaf conductance measurements and associated data analyses under the supervision of R.B. and S.D.. P.G. and N.F. performed stem sampling. A.M., N.F. and P.G. were responsible for cutting, staining and imaging stem cross-sections, with technical support from S.M.. I.S. and P.G. prepared samples and performed SEM observations, with scientific support from F.L.. A.M., P.G. and N.F. analysed histological scans, with scientific support from S.M. and F.L.. A.M. and P.G. prepared the samples for metabolomics. C.R., P.P. and the MetaboHUB-Bordeaux team performed untargeted metabolomic experiments (extraction, LC-MS and MS-DIAL analysis). P.P. contributed to metabolomic data analysis and interpretation. P.G. and N.F. ground the samples and performed DNA extraction, and PCR for metabarcoding. P.G. analysed all datasets, produced the figures and wrote the first version of the paper under the supervision of C.E.L.D. All the authors contributed to scientific discussions, critically revised and approved the final version of the manuscript.

## Data availability

All sequencing datasets (ITS and 16S metabarcoding) are under the process of being made available from the ENA database under accession number PRJEB100545. The raw datasets (ecophysiology, wood anatomy, metabolism) will be made available *via* data.gouv.fr upon publication of the manuscript. The code used to compute minimum leaf conductance is available from the following Gitlab repository: https://gitub.u-bordeaux.fr/phenobois.

## Supporting information

**Supplementary Table S1**: List of the 46 *Vitis vinifera* L. cultivars studied.

**Supplementary Table S2**: List of metabolic features differentially abundant between stem health statuses and varietal susceptibility clusters.

**Supplementary Table S3**: Effect of cultivar on *g_s_* and *g_min_*.

**Supplementary Table S4**: Effects of cultivar, vascular bundle type, their interaction and stem health status on stem wood anatomical traits.

**Supplementary Table S5**: Effects of cultivar, health status and/or the presence of vascular occlusions on grapevine physiological responses to esca.

**Supplementary Table S6**: OTU table corresponding to the entire decontaminated ITS metabarcoding dataset.

**Supplementary Table S7**: OTU table corresponding to the entire decontaminated 16S metabarcoding dataset.

**Supplementary Table S8**: Effects of stem health status, varietal susceptibility cluster, cultivar and/or esca symptom severity on Shannon index and the beta-diversity of stem microbial communities.

**Supplementary Figure S1. Varietal gradients of leaf gas exchange traits.** (A) Mean ± SEM of stomatal conductance measured in June 2024 on 46 cultivars (*n* = 1,840 plants) (B) Mean ± SEM of minimum leaf conductance measured in May 2024 on detached leaves from 19 cultivars (*n* = 114 leaves). Bars are coloured according to varietal mean esca susceptibility (mean foliar symptom incidence from 2017 to 2024). *P*-values correspond to cultivar effects in linear mixed models.

**Supplementary Figure S2. Variability of stem xylem anatomical traits among grapevine cultivars and relationship to varietal esca susceptibility.** (A) Boxplots representing the mean number of xylem ray parenchyma layers for 10 cultivars (*n* = 60 stems). (B) Boxplots representing mean xylem fibre lumen area for 10 cultivars (*n* = 60 stems). (C) Boxplots representing intervessel pit density across 13 cultivars (*n* = 42 stems). Boxplots displaying the median and interquartile range, with whiskers extending to the minimum and maximum values, excluding outliers, which are shown as individual black points. The plots are coloured according to varietal mean esca susceptibility (mean foliar symptom incidence from 2017 to 2024). *P*-values correspond to cultivar effects in linear mixed models. The letters correspond to the significance groups in Tukey tests with an alpha risk of 5%.

**Supplementary Figure S3. Co-occurrence of vascular occlusions and starch storage in esca/tiger-stripe stems (*n* = 9 cultivars; *n* = 25 stems)**. Boxplots represent the percentage of the ray parenchyma area storing starch in esca/tiger-stripe stems according to the presence of vascular occlusions in the adjacent vascular bundles. Boxplots display the median and interquartile range, with whiskers extending to the minimum and maximum values, excluding outliers, which are shown as individual black points. The plots are coloured according to the presence of vascular occlusions. *P*-values correspond to the effect of vascular occlusions in a linear mixed model. The letters correspond to the significance groups in Tukey tests with an alpha risk of 5%.

**Supplementary Figure S4. Cultivar effect on the co-occurrence of vascular occlusions and starch storage in esca/tiger-stripe stems (*n* = 9 cultivars; *n* = 25 stems).** Boxplots represent the percentage of the ray parenchyma area storing starch in esca/tiger-stripe stems according to the presence of vascular occlusions in the adjacent vascular bundles. Boxplots display the median and interquartile range, with whiskers extending to the minimum and maximum values, excluding outliers, which are shown as individual black points. The plots are coloured according to the presence of vascular occlusions. Panels are ordered according to varietal mean esca susceptibility.

**Supplementary Figure S5. Effects of esca expression on leaf gas exchange across five cultivars.** (A, C) Boxplots representing maximal leaf stomatal conductance (A) before (*n* = 34 plants) and (C) after esca foliar symptom expression (*n* = 37 plants) on control plants and plants displaying esca symptoms during the season. (B, D) Boxplots representing maximal leaf carbon assimilation (B) before (*n* = 34 plants) and (D) after esca foliar symptom expression (*n* = 37 plants) on control plants and plants displaying esca symptoms during the season. Boxplots are coloured according to esca expression. *P*-values correspond to the effect of stem health status in a linear mixed model. The letters correspond to significance groups in Tukey tests with an alpha risk of 5%.

**Supplementary Figure S6. Distribution of the 20 most abundant fungal species in *Vitis vinifera* stems.** (A) Distribution between stem health statuses. (B) Distribution between varietal esca susceptibility classes in control stems. Mean relative abundances were calculated for each species. Stacked bars are coloured according to the OTU. Less abundant species are grouped in the category “Other” and shown in grey.

**Supplementary Figure S7. Distribution of the 20 most abundant bacterial genera in *Vitis vinifera* stems.** (A) Distribution between stem health statuses. (B) Distribution between varietal esca susceptibility classes in control stems. Mean relative abundances were calculated for each genus. Stacked bars are coloured according to the OTU. Less abundant genera are grouped in the category “Other” and shown in grey.

**Supplementary Figure S8. Effects of varietal esca susceptibility on control stem microbial communities** (*n* = 10 cultivars, *n* = 29 stems). (A, B) Effect of esca varietal susceptibility class on the Shannon index associated with (A) fungal and (B) bacterial communities. Boxplots display the median and interquartile range, with whiskers extending to the minimum and maximum values, excluding outliers, which are shown as individual black points. (C, D) PCA summarising the effects of esca varietal susceptibility class on the structure of (C) fungal and (D) bacterial communities, determined with CLR-transformed data. Ellipses correspond to the 95% confidence interval for each group. The colours and shapes of the points indicate esca varietal susceptibility class. *P*-values correspond to esca effects in ANOVA on Shannon index, or PERMANOVA.

**Supplementary Figure S9. Effects of esca symptom severity on esca/tiger-stripe stem microbial communities** (*n* = 9 cultivars, *n* = 29 stems). (A, C) Effect of esca symptom severity on the Shannon index associated with (A) fungal and (C) bacterial communities. Boxplots display the median and interquartile range, with whiskers extending to the minimum and maximum values, excluding outliers, which are shown as individual black points. (B, D) PCA summarising the effects of esca symptom severity on the structure of (B) fungal and (D) bacterial communities performed with CLR-transformed data. Ellipses correspond to the 95% confidence interval for each group. The colours and shapes of all points correspond to esca symptom severity. *P*-values correspond to the effects of esca in ANOVA on Shannon index, or PERMANOVA.

**Supplementary Figure S10. Varietal relationships between pit-related traits and esca foliar symptom incidence in summer 2024.** (A) Mean varietal intervessel pit density on 13 cultivars. (B) Mean varietal aperture fraction of intervessel pits on 13 cultivars. Each dot corresponds to the value for a specific cultivar measured. Dots are colored according to varietal average esca susceptibility (mean foliar symptom incidence from 2017 to 2024). Correlation coefficients and associated p-values are computed in Pearson’s tests.

**Supplementary Figure S11. Effects of varietal esca susceptibility on the metabolome in control stems (*n* = 10 cultivars, *n* = 29 stems).** (A) PCA summarising the effects of varietal esca susceptibility on the structure of the control stem metabolome determined with normalised, transformed and scaled data. Ellipses correspond to the 95% confidence interval for each group. (B, C, D) Volcano plots for all features. The cut-off values for significance are set at |log_2_FC| > 2 and *P* < 0.01. Non-significant (NS) and putatively contaminant features are coloured in grey. (E, F, G) Class assignment of differentially abundant features selected in Volcano analyses (cut-off values set at |log_2_FC| > 2 and *P* < 0.01). Class assignment was performed with Classyfire and published data. Features that could not be assigned to any known compound or metabolic class are classified as “Unknown”. Metabolic classes were classified as belonging to primary (I, above) or secondary (II, below) metabolism. The colours and shapes of all dots and ellipses correspond to esca varietal susceptibility class. *P*-values correspond to the effect of varietal esca susceptibility in PERMANOVA.

**Supplementary Figure S12. Effects of plant esca expression on the asymptomatic stem metabolome and relationship to esca varietal susceptibility.** Panels are grouped by esca susceptibility: (A, D, G) weakly susceptible cultivars (*n* = 3 cultivars, *n* = 12 stems), (B, E, H) moderately susceptible cultivars (*n* = 3 cultivars, *n* = 21 stems), and (C, F, I) highly susceptible cultivars (*n* = 4 cultivars, *n* = 29 stems). (A, B, C) PCA summarising the effects of esca expression on the structure of the stem metabolome performed on normalised, transformed and scaled data. Ellipses correspond to the 95% confidence interval for each group. Dots and ellipses are coloured and shaped according to stem health status. (D, F, E) Volcano plots for all features. The cut-off values for significance are set at |log_2_FC| > 2 and *P* < 0.01. Non-significant (NS) and putative contaminant features are shown in grey. Features for which enrichment was observed in control stems are shown as blue circles, and features for which enrichment was observed in esca/asymptomatic stems are shown as purple diamonds. (G, H, I) Class assignment of differentially abundant features selected in Volcano analyses (cut-off values set at |log_2_FC| > 2 and *P* < 0.01). Class assignment was performed with Classyfire and published data. Classes for which enrichment was observed in control stems are shown as blue circles, and classes for which enrichment was observed in esca/asymptomatic stems are shown as purple diamonds. Features that could not be assigned to any known compound or metabolic class are classified as “Unknown”. Metabolic classes were classified as belonging to primary (I, above) or secondary (II, below) metabolism.

**Supplementary Figure S13. Effects of stem health status on the stem metabolome in symptomatic plants and relationship to esca varietal susceptibility.** Panels are grouped by esca susceptibility: (A, D, G) weakly susceptible cultivars (*n* = 3 cultivars, *n* = 7 stems), (B, E, H) moderately susceptible cultivars (*n* = 3 cultivars, *n* = 22 stems), and (C, F, I) highly susceptible cultivars (*n* = 4 cultivars, *n* = 33 stems). (A, B, C) PCA summarising the effects of esca expression the structure of the stem metabolome performed on normalised, transformed and scaled data. Ellipses correspond to the 95% confidence interval for each group. Dots and ellipses are coloured and shaped according to stem health status. (D, F, E) Volcano plots for all features. The cut-off values for significance are set at |log_2_FC| > 2 and *P* < 0.01. Non-significant (NS) and putative contaminant features are shown in grey. Features for which enrichment was observed in esca/asymptomatic stems are shown as purple diamonds, and features for which enrichment was observed in esca/tiger-stripe stems are shown as yellow triangles. (G, H, I). Class assignment of differentially abundant features selected in Volcano analyses (cut-off values set at |log_2_FC| > 2 and *P* < 0.01). Class assignment was performed with Classyfire and published data. Classes for which enrichment was observed in esca/asymptomatic stems are shown as purple diamonds, and features for which enrichment was observed in esca/tiger-stripe stems are shown as yellow triangles. Features that could not be assigned to any known compound or metabolic class are classified as “Unknown”. Metabolic classes were classified as belonging to primary (I, above) or secondary (II, below) metabolism.

**Supplementary Figure S14. Effects of esca symptom severity on the metabolome in esca/tiger-stripe stems (*n* = 9 cultivars, *n* = 29 stems).** (A) PCA summarising the effects of esca symptom severity on the structure of the esca/tiger-stripe stem metabolome performed on normalised, transformed and scaled data. Ellipses correspond to the 95% confidence interval for each group. (B, C, D) Volcano plots for all features. The cut-off values for significance are set at |log_2_FC| > 2 and *P* < 0.01. Non-significant (NS) and putative contaminant features are shown in grey. (E, F, G) Class assignment of differentially abundant features selected in Volcano analyses (cut-off values set at |log_2_FC| > 2 and *P* < 0.01). Class assignment was performed with Classyfire and published data. Features that could not be assigned to any known compound or metabolic class are classified as “Unknown”. Metabolic classes were classified as belonging to primary (I, above) or secondary (II, below) metabolism. All dots and ellipses are coloured and shaped according to esca symptom severity. *P*-values correspond to the effect of esca symptom severity in PERMANOVA.

**Supplementary Figure S15. Varietal relationships between pit-related traits and esca foliar symptom incidence in summer 2024.** (A) Mean varietal intervessel pit density on 13 cultivars. (B) Mean varietal aperture fraction of intervessel pits on 13 cultivars. Each dot corresponds to the value for a specific cultivar measured. Dots are colored according to varietal average esca susceptibility (mean foliar symptom incidence from 2017 to 2024). Correlation coefficients and associated p-values are computed in Pearson’s tests.

## Notes

### Competing Interest Statement

The authors have declared no competing interest.

### Summary of Updates

Minor corrections have been made to the following: the title and abstract; the description of the methods to make them more concise; and various minor changes in the body of the text.

