## Supplementary figures for "Intraspecific variation in functional strategies underlies susceptibility to a vascular disease in grapevine"

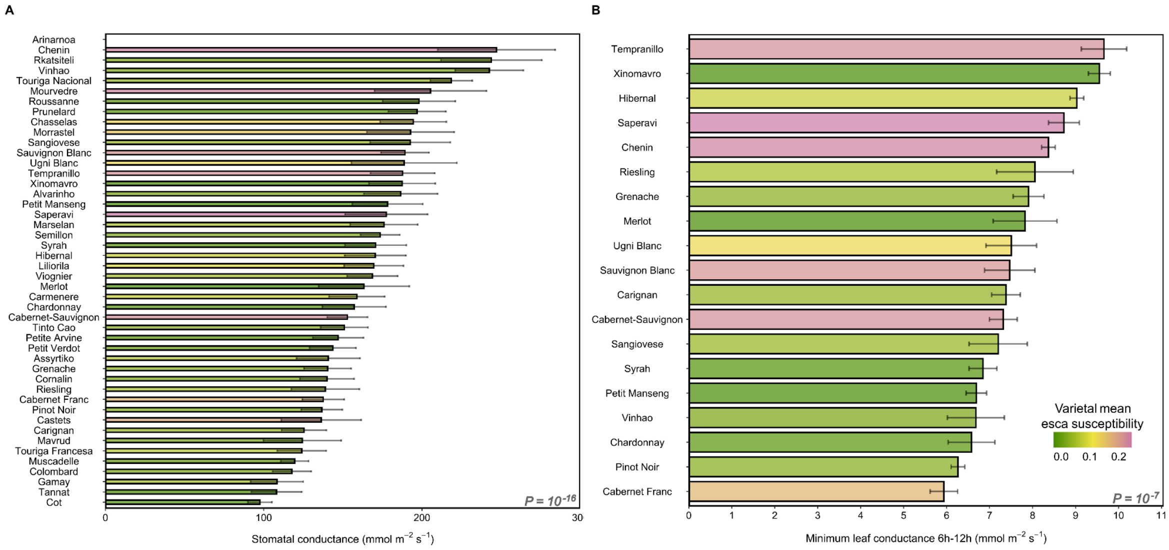


**Supplementary Figure S1. Varietal gradients of leaf gas exchange traits.** (A) Mean ± SEM of stomatal conductance measured in June 2024 on 46 cultivars (*n* = 1,840 plants) (B) Mean ± SEM of minimum leaf conductance measured in May 2024 on detached leaves from 19 cultivars (*n* = 114 leaves). Bars are coloured according to varietal mean esca susceptibility (mean foliar symptom incidence from 2017 to 2024). *P*-values correspond to cultivar effects in linear mixed models.

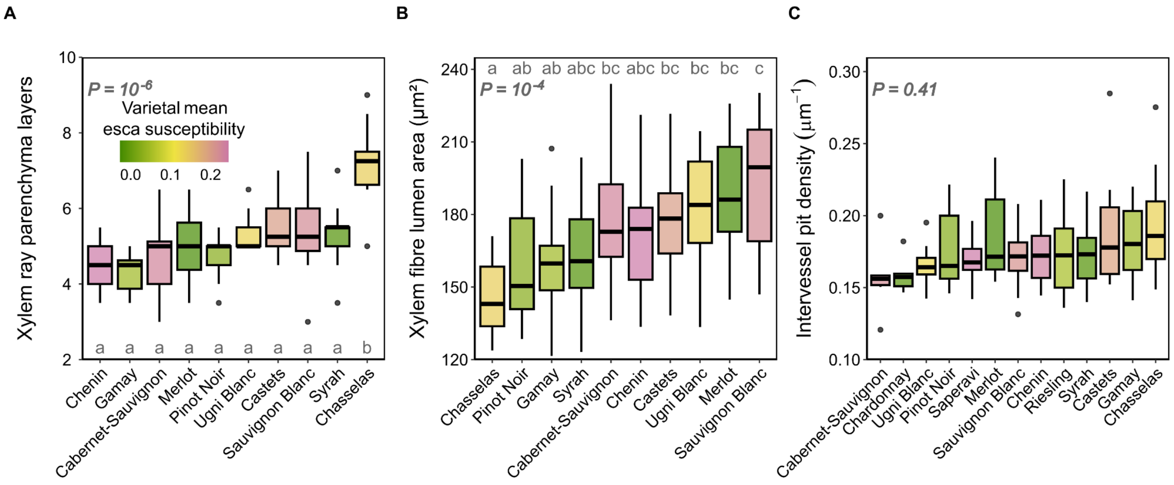


**Supplementary Figure S2. Variability of stem xylem anatomical traits among grapevine cultivars and relationship to varietal esca susceptibility.** (A) Boxplots representing the mean number of xylem ray parenchyma layers for 10 cultivars (*n* = 60 stems). (B) Boxplots representing mean xylem fibre lumen area for 10 cultivars (*n* = 60 stems). (C) Boxplots representing intervessel pit density across 13 cultivars (*n* = 42 stems). Boxplots displaying the median and interquartile range, with whiskers extending to the minimum and maximum values, excluding outliers, which are shown as individual black points. The plots are coloured according to varietal mean esca susceptibility (mean foliar symptom incidence from 2017 to 2024). *P*-values correspond to cultivar effects in linear mixed models. The letters correspond to the significance groups in Tukey tests with an alpha risk of 5%.

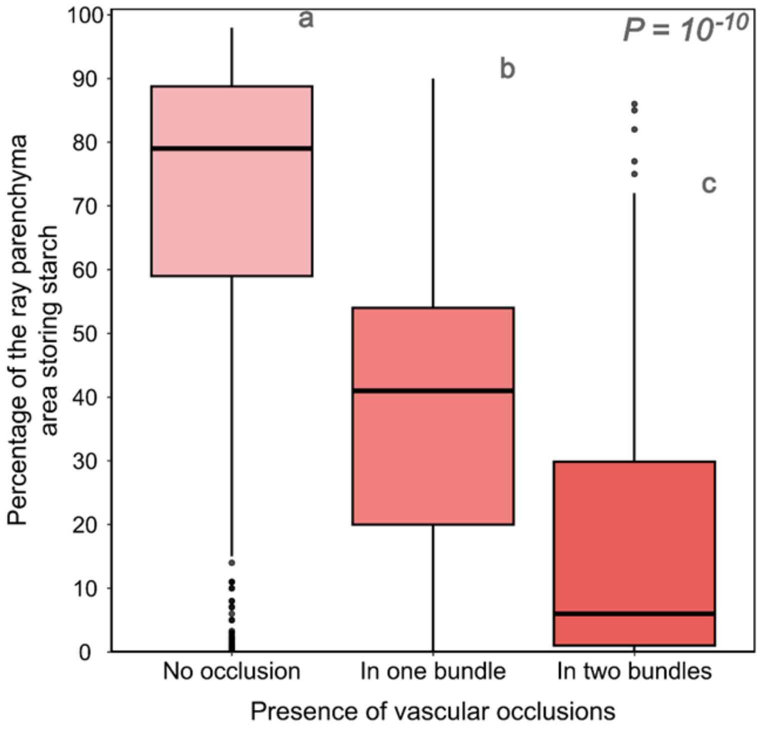


**Supplementary Figure S3. Co-occurrence of vascular occlusions and starch storage in esca/tiger-stripe stems (*n* = 9 cultivars; *n* = 25 stems).** Boxplots represent the percentage of the ray parenchyma area storing starch in esca/tiger-stripe stems according to the presence of vascular occlusions in the adjacent vascular bundles. Boxplots display the median and interquartile range, with whiskers extending to the minimum and maximum values, excluding outliers, which are shown as individual black points. The plots are coloured according to the presence of vascular occlusions. *P*-values correspond to the effect of vascular occlusions in a linear mixed model. The letters correspond to the significance groups in Tukey tests with an alpha risk of 5%.

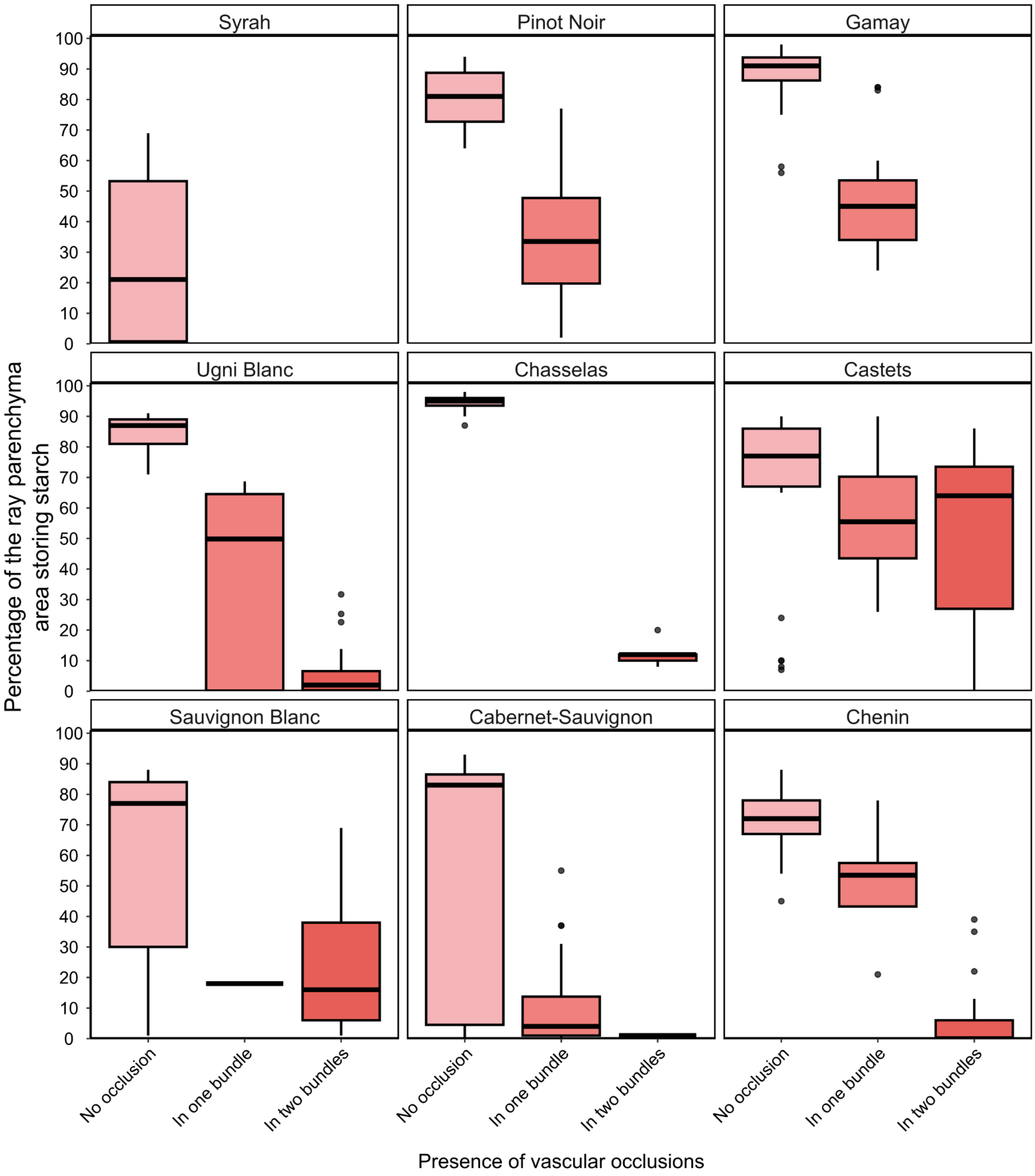


**Supplementary Figure S4. Cultivar effect on the co-occurrence of vascular occlusions and starch storage in esca/tiger-stripe stems (*n* = 9 cultivars; *n* = 25 stems).** Boxplots represent the percentage of the ray parenchyma area storing starch in esca/tiger-stripe stems according to the presence of vascular occlusions in the adjacent vascular bundles. Boxplots display the median and interquartile range, with whiskers extending to the minimum and maximum values, excluding outliers, which are shown as individual black points. The plots are coloured according to the presence of vascular occlusions. Panels are ordered according to varietal mean esca susceptibility.

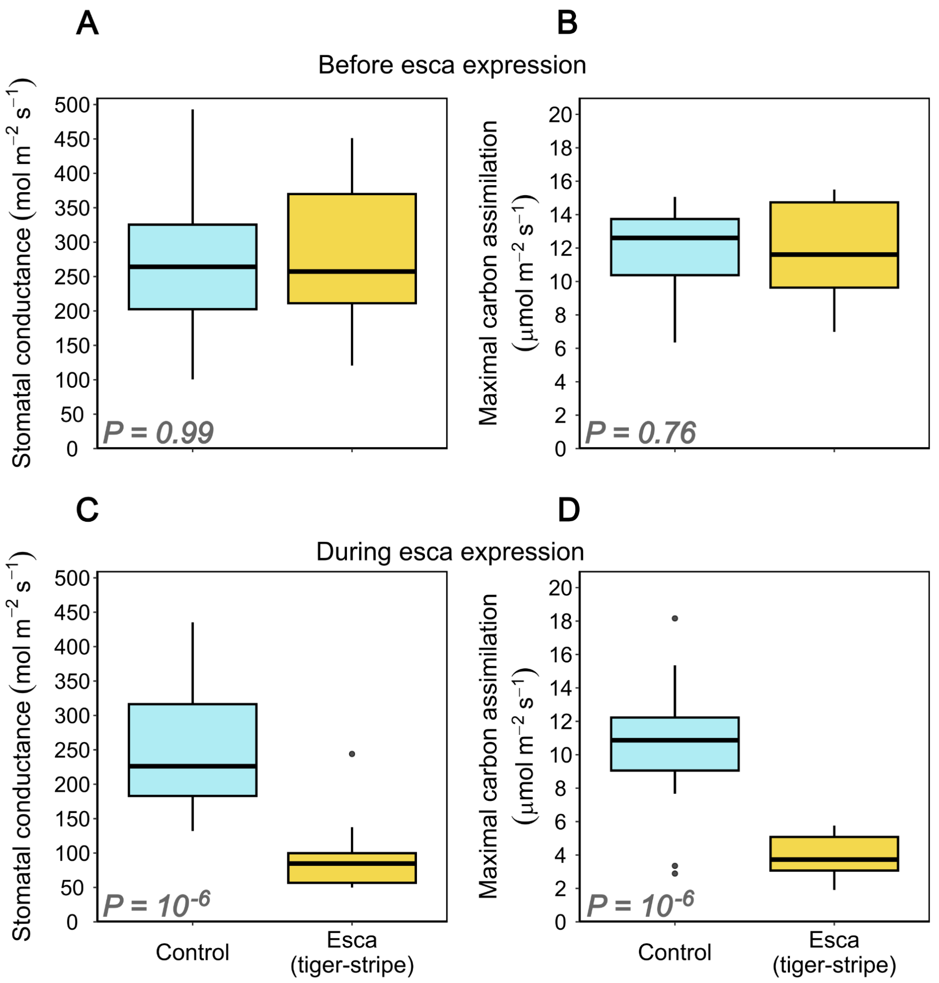


**Supplementary Figure S5. Effects of esca expression on leaf gas exchange across five cultivars.** (A, C) Boxplots representing maximal leaf stomatal conductance (A) before ($n$ = 34 plants) and (C) after esca foliar symptom expression ($n$ = 37 plants) on control plants and plants displaying esca symptoms during the season. (B, D) Boxplots representing maximal leaf carbon assimilation (B) before ($n$ = 34 plants) and (D) after esca foliar symptom expression ($n$ = 37 plants) on control plants and plants displaying esca symptoms during the season. Boxplots are coloured according to esca expression. *P*-values correspond to the effect of stem health status in a linear mixed model. The letters correspond to significance groups in Tukey tests with an alpha risk of 5%.

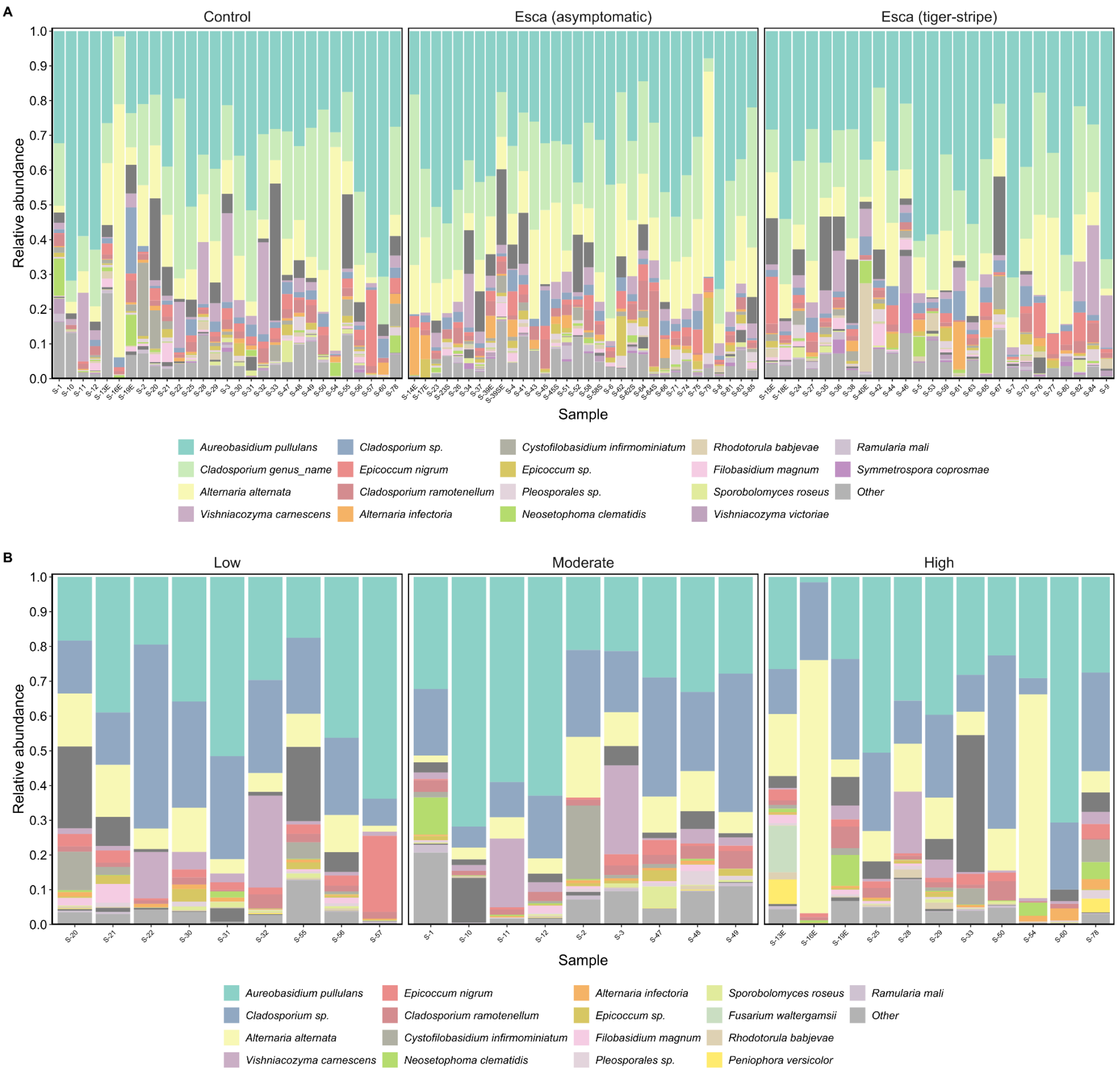


**Supplementary Figure S6. Distribution of the 20 most abundant fungal species in *Vitis vinifera* stems.** (A) Distribution between stem health statuses. (B) Distribution between varietal esca susceptibility classes in control stems. Mean relative abundances were calculated for each species. Stacked bars are coloured according to the OTU. Less abundant species are grouped in the category “Other” and shown in grey.

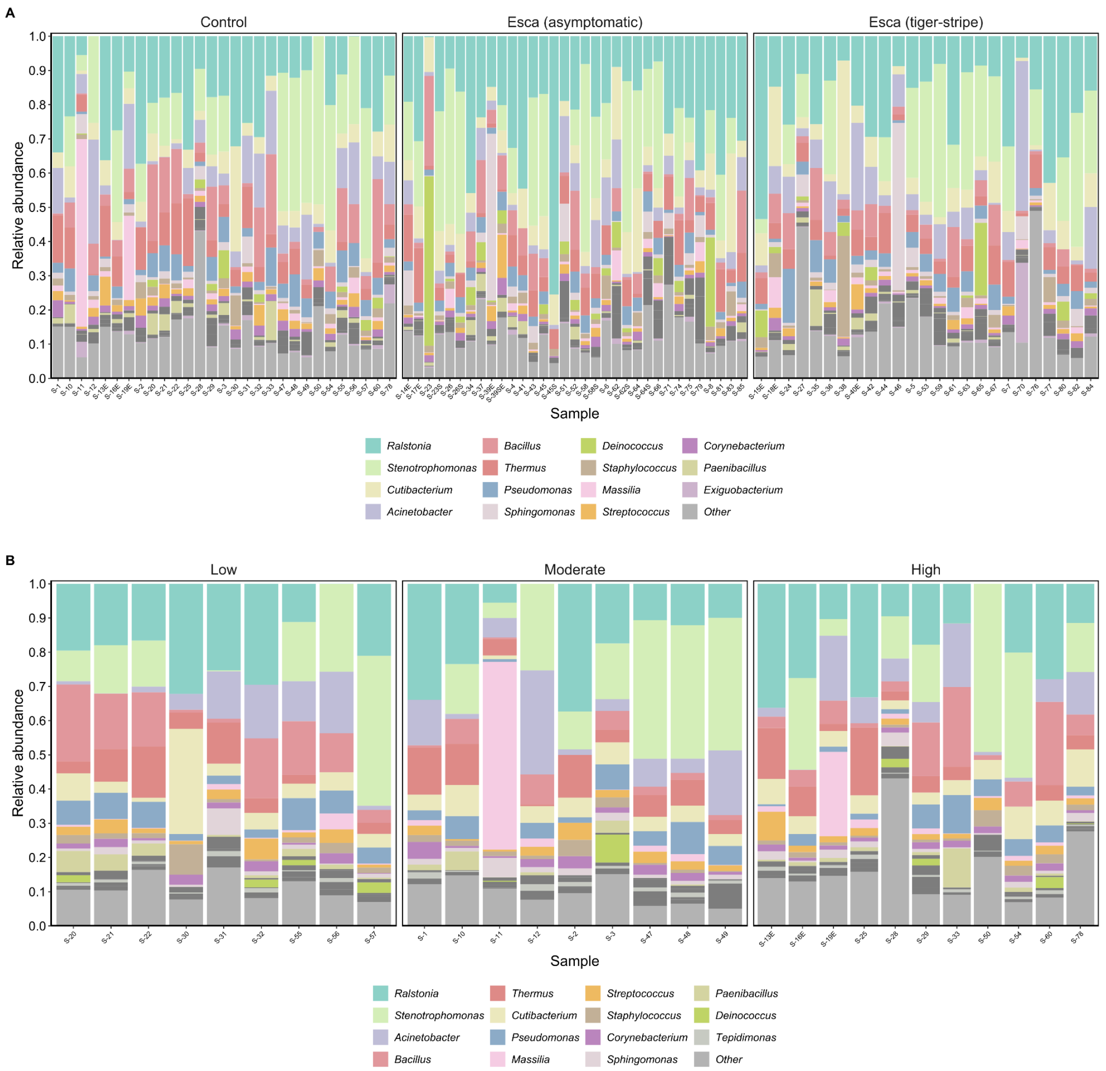


**Supplementary Figure S7. Distribution of the 20 most abundant bacterial genera in *Vitis vinifera* stems.** (A) Distribution between stem health statuses. (B) Distribution between varietal esca susceptibility classes in control stems. Mean relative abundances were calculated for each genus. Stacked bars are coloured according to the OTU. Less abundant genera are grouped in the category “Other” and shown in grey.

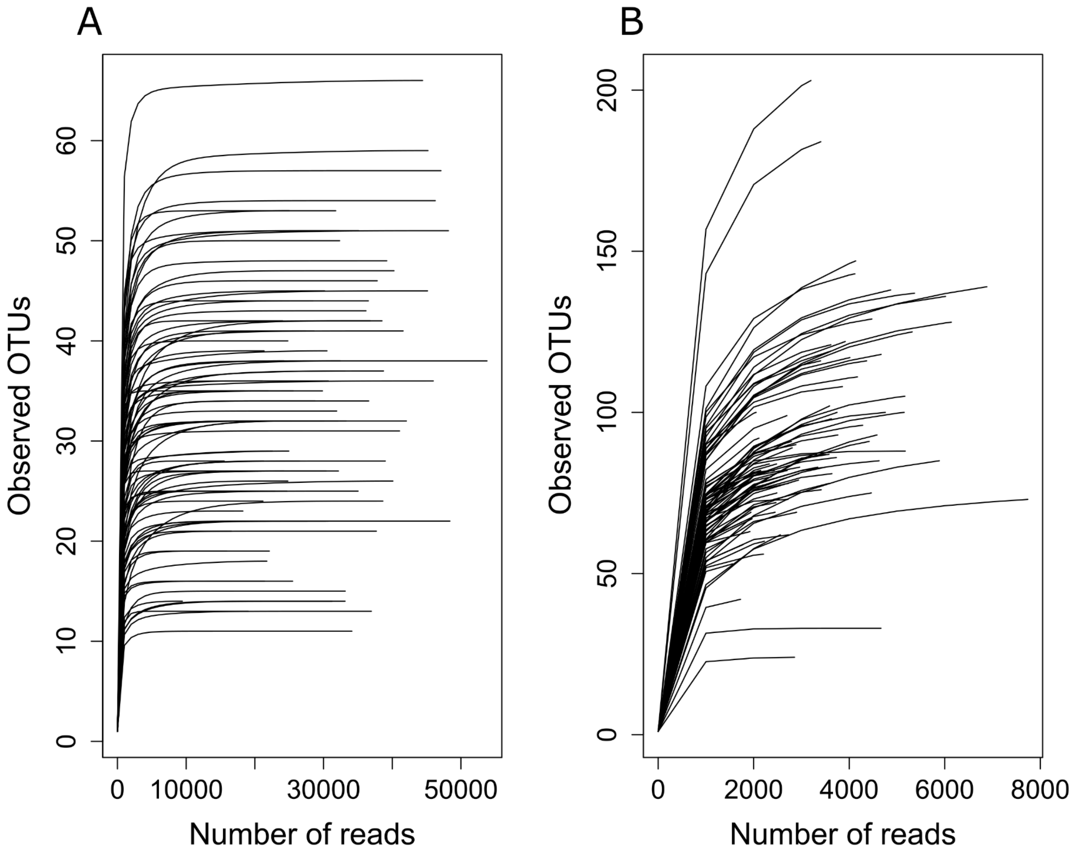


**Supplementary Fig. S8. Rarefaction curves of grapevine stem microbial communities (fungi and bacteria).** Rarefaction curves showing observed OTU richness as a function of sequencing depth for **(A)** ITS and **(B)** 16S datasets.

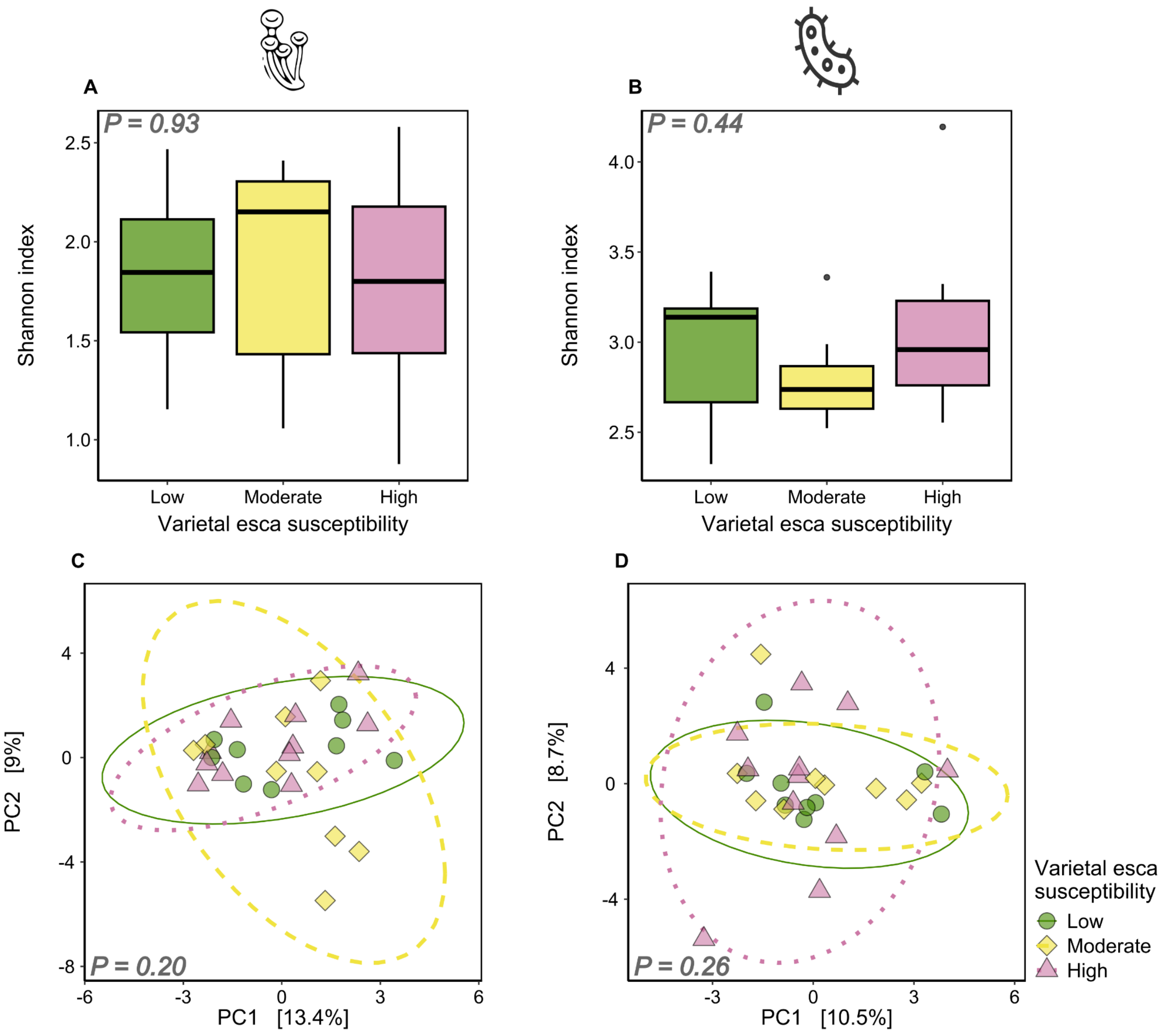


**Supplementary Figure S9. Effects of varietal esca susceptibility on control stem microbial communities** (*n* = 10 cultivars, *n* = 29 stems). (A, B) Effect of esca varietal susceptibility class on the Shannon index associated with (A) fungal and (B) bacterial communities. Boxplots display the median and interquartile range, with whiskers extending to the minimum and maximum values, excluding outliers, which are shown as individual black points. (C, D) PCA summarising the effects of esca varietal susceptibility class on the structure of (C) fungal and (D) bacterial communities, determined with CLR-transformed data. Ellipses correspond to the 95% confidence interval for each group. The colours and shapes of the points indicate esca varietal susceptibility class. *P*-values correspond to esca effects in ANOVA on Shannon index, or PERMANOVA.

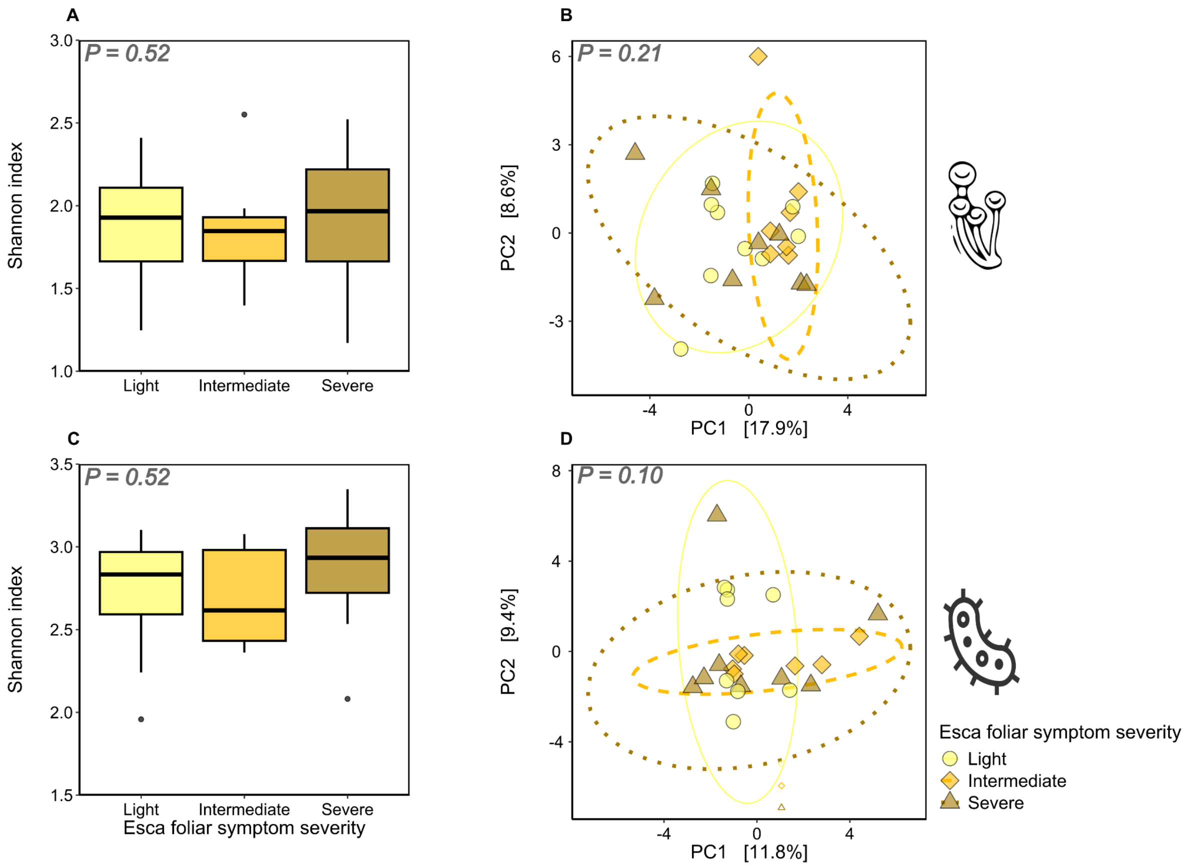


**Supplementary Figure S10. Effects of esca symptom severity on esca/tiger-stripe stem microbial communities** ($n$ = 9 cultivars, $n$ = 29 stems). (A, C) Effect of esca symptom severity on the Shannon index associated with (A) fungal and (C) bacterial communities. Boxplots display the median and interquartile range, with whiskers extending to the minimum and maximum values, excluding outliers, which are shown as individual black points. (B, D) PCA summarising the effects of esca symptom severity on the structure of (B) fungal and (D) bacterial communities performed with CLR-transformed data. Ellipses correspond to the 95% confidence interval for each group. The colours and shapes of all points correspond to esca symptom severity. *P*-values correspond to the effects of esca in ANOVA on Shannon index, or PERMANOVA.

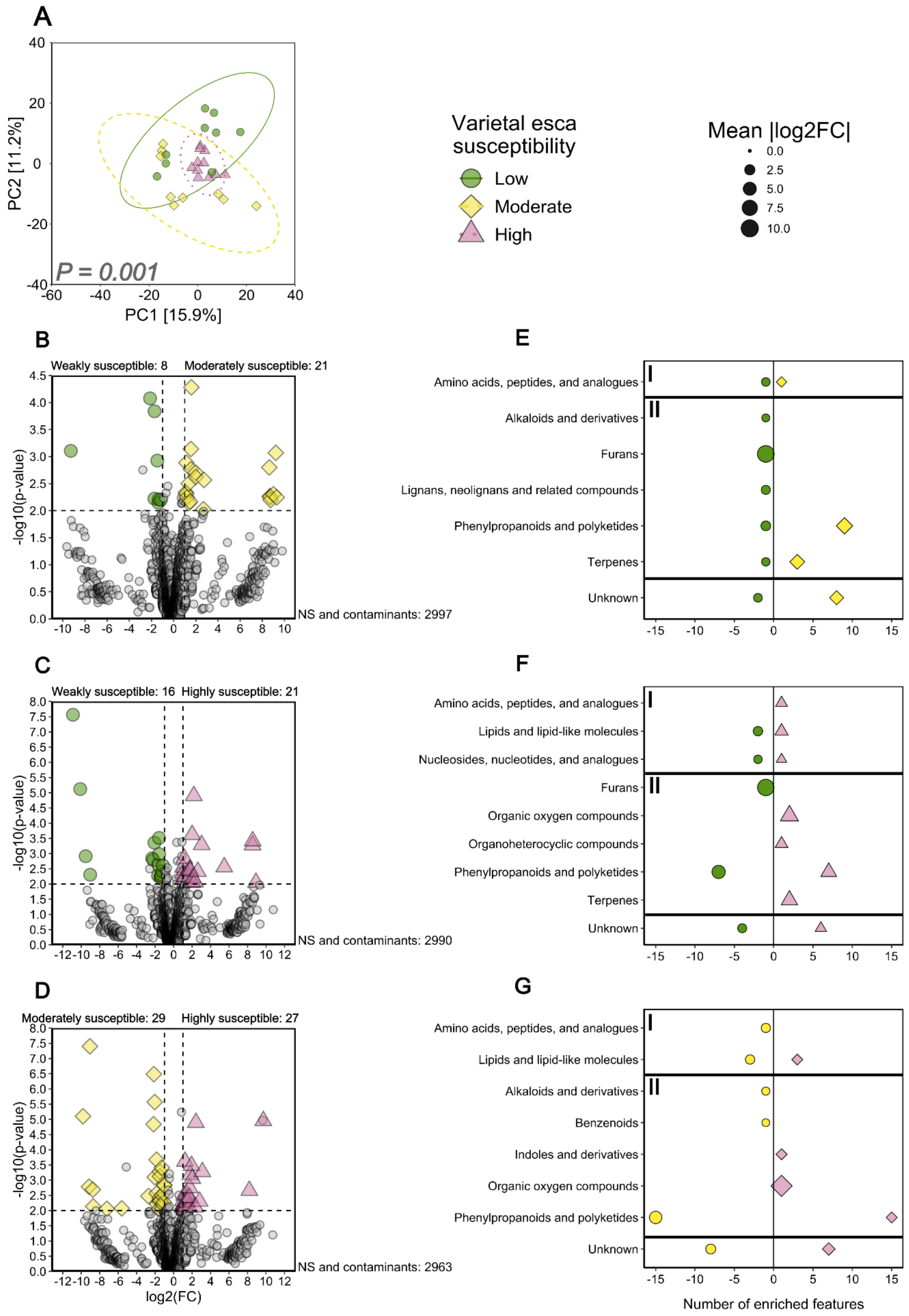


**Supplementary Figure S11. Effects of varietal esca susceptibility on the metabolome in control stems (*n* = 10 cultivars, *n* = 29 stems).** (A) PCA summarising the effects of varietal esca susceptibility on the structure of the control stem metabolome determined with normalised, transformed and scaled data. Ellipses correspond to the 95% confidence interval for each group. (B, C, D) Volcano plots for all features. The cut-off values for significance are set at $|log_2FC| > 2$ and $P < 0.01$. Non-significant (NS) and putatively contaminant features are coloured in grey. (E, F, G) Class assignment of differentially abundant features selected in Volcano analyses (cut-off values set at $|log_2FC| > 2$ and $P < 0.01$). Class assignment was performed with Classyfire and published data. Features that could not be assigned to any known compound or metabolic class are classified as “Unknown”. Metabolic classes were classified as belonging to primary (I, above) or secondary (II, below) metabolism. The colours and shapes of all dots and ellipses correspond to esca varietal susceptibility class. *P*-values correspond to the effect of varietal esca susceptibility in PERMANOVA.

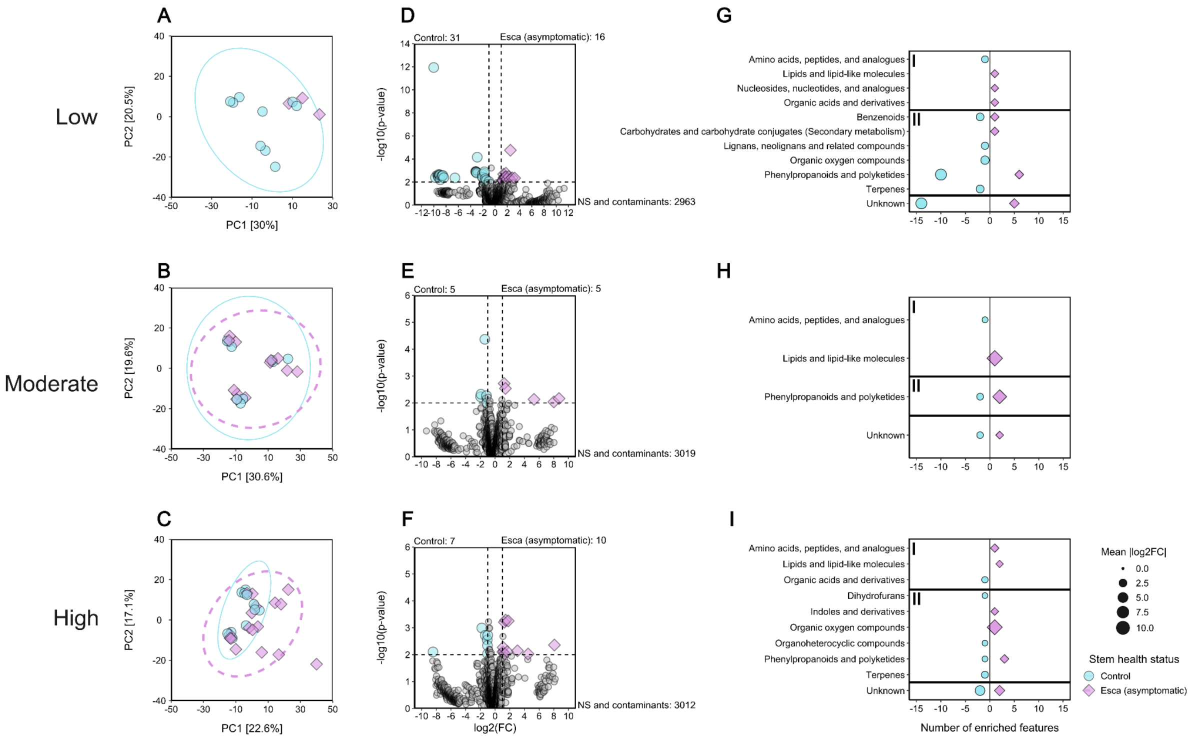


**Supplementary Figure S12. Effects of plant esca expression on the asymptomatic stem metabolome and relationship to esca varietal susceptibility.** Panels are grouped by esca susceptibility: (A, D, G) weakly susceptible cultivars (*n* = 3 cultivars, *n* = 12 stems), (B, E, H) moderately susceptible cultivars (*n* = 3 cultivars, *n* = 21 stems), and (C, F, I) highly susceptible cultivars (*n* = 4 cultivars, *n* = 29 stems). (A, B, C) PCA summarising the effects of esca expression on the structure of the stem metabolome performed on normalised, transformed and scaled data. Ellipses correspond to the 95% confidence interval for each group. Dots and ellipses are coloured and shaped according to stem health status. (D, F, E) Volcano plots for all features. The cut-off values for significance are set at $|\log_2 FC| > 2$ and $P < 0.01$. Non-significant (NS) and putative contaminant features are shown in grey. Features for which enrichment was observed in control stems are shown as blue circles, and features for which enrichment was observed in esca/asymptomatic stems are shown as purple diamonds. (G, H, I) Class assignment of differentially abundant features selected in Volcano analyses (cut-off values set at $|\log_2 FC| > 2$ and $P < 0.01$). Class assignment was performed with Classyfire and published data. Classes for which enrichment was observed in control stems are shown as blue circles, and classes for which enrichment was observed in esca/asymptomatic stems are shown as purple diamonds. Features that could not be assigned to any known compound or metabolic class are classified as “Unknown”. Metabolic classes were classified as belonging to primary (I, above) or secondary (II, below) metabolism.

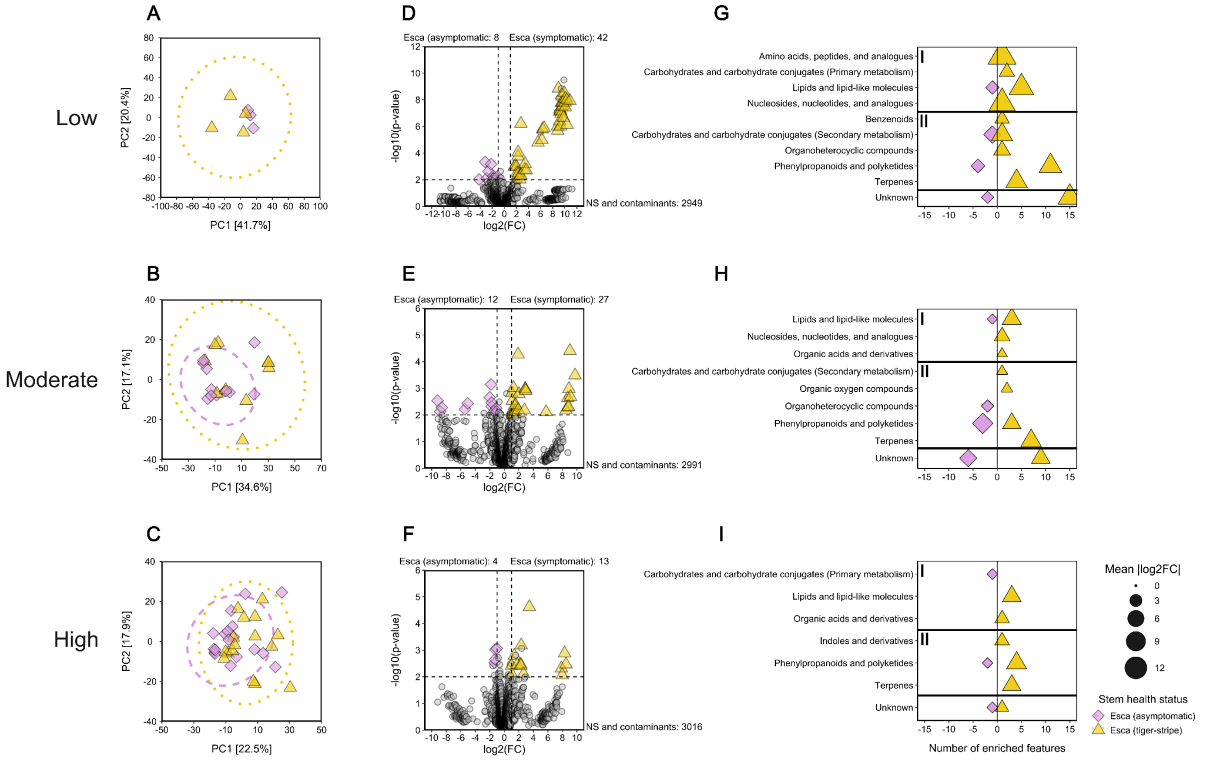


**Supplementary Figure S13. Effects of stem health status on the stem metabolome in symptomatic plants and relationship to esca varietal susceptibility.** Panels are grouped by esca susceptibility: (A, D, G) weakly susceptible cultivars ($n$ = 3 cultivars, $n$ = 7 stems), (B, E, H) moderately susceptible cultivars ($n$ = 3 cultivars, $n$ = 22 stems), and (C, F, I) highly susceptible cultivars ($n$ = 4 cultivars, $n$ = 33 stems). (A, B, C) PCA summarising the effects of esca expression the structure of the stem metabolome performed on normalised, transformed and scaled data. Ellipses correspond to the 95% confidence interval for each group. Dots and ellipses are coloured and shaped according to stem health status. (D, F, E) Volcano plots for all features. The cut-off values for significance are set at $|\log_2 FC| > 2$ and $P < 0.01$. Non-significant (NS) and putative contaminant features are shown in grey. Features for which enrichment was observed in esca/asymptomatic stems are shown as purple diamonds, and features for which enrichment was observed in esca/tiger-stripe stems are shown as yellow triangles. (G, H, I). Class assignment of differentially abundant features selected in Volcano analyses (cut-off values set at $|\log_2 FC| > 2$ and $P < 0.01$). Class assignment was performed with Classyfire and published data. Classes for which enrichment was observed in esca/asymptomatic stems are shown as purple diamonds, and features for which enrichment was observed in esca/tiger-stripe stems are shown as yellow triangles. Features that could not be assigned to any known compound or metabolic class are classified as “Unknown”. Metabolic classes were classified as belonging to primary (I, above) or secondary (II, below) metabolism.

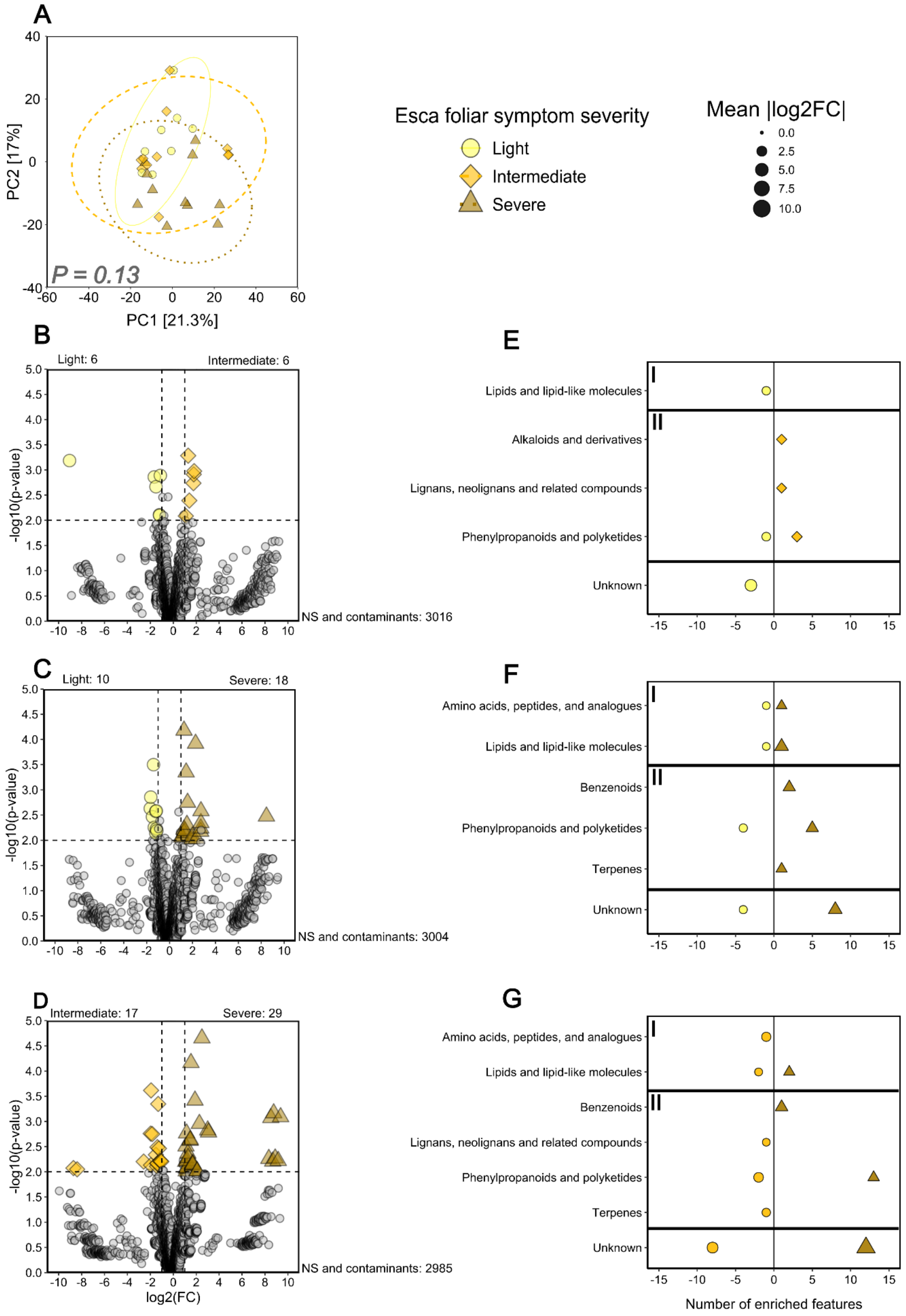


**Supplementary Figure S14. Effects of esca symptom severity on the metabolome in esca/tiger-stripe stems (*n* = 9 cultivars, *n* = 29 stems).** (A) PCA summarising the effects of esca symptom severity on the structure of the esca/tiger-stripe stem metabolome performed on normalised, transformed and scaled data. Ellipses correspond to the 95% confidence interval for each group. (B, C, D) Volcano plots for all features. The cut-off values for significance are set at $|log_2FC| > 2$ and $P < 0.01$. Non-significant (NS) and putative contaminant features are shown in grey. (E, F, G) Class assignment of differentially abundant features selected in Volcano analyses (cut-off values set at $|log_2FC| > 2$ and $P < 0.01$). Class assignment was performed with Classyfire and published data. Features that could not be assigned to any known compound or metabolic class are classified as “Unknown”. Metabolic classes were classified as belonging to primary (I, above) or secondary (II, below) metabolism. All dots and ellipses are coloured and shaped according to esca symptom severity. *P*-values correspond to the effect of esca symptom severity in PERMANOVA.

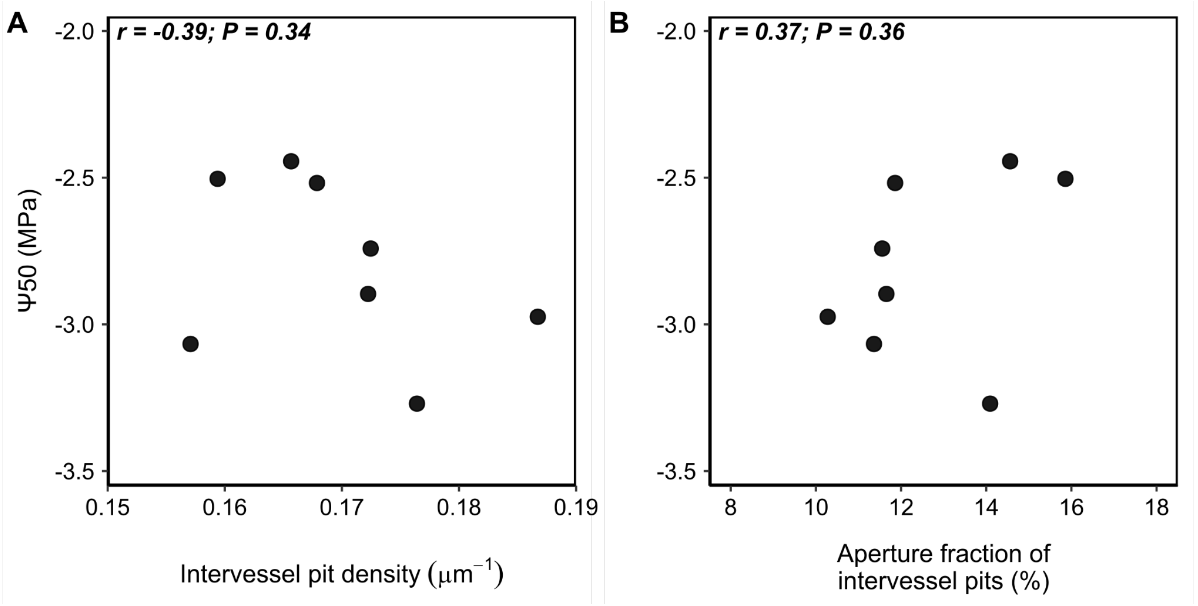


**Supplementary Figure S15. Varietal relationships between pit-related traits and esca foliar symptom incidence in summer 2024.** (A) Mean varietal intervessel pit density on 13 cultivars. (B) Mean varietal aperture fraction of intervessel pits on 13 cultivars. Each dot corresponds to the value for a specific cultivar measured. Dots are colored according to varietal average esca susceptibility (mean foliar symptom incidence from 2017 to 2024). Correlation coefficients and associated p-values are computed in Pearson's tests.
